# Variability in Forest Plant Traits along the Western Ghats of India and Their Environmental Drivers at Different Resolutions

**DOI:** 10.1101/2023.02.01.526666

**Authors:** Ting Zheng, Aditya Singh, Ankur R. Desai, N.K. Krishnayya, Philip A. Townsend

## Abstract

Identifying key environmental drivers for plant functional traits is an important step to understanding and predicting ecosystem responses to a changing climate. Imaging spectroscopy offers great potential to map plant traits at fine resolution across broad regions and then assess controls on their variation across spatial resolutions. We applied permutational partial least-squares regression to map seven key foliar chemical and morphological traits using NASA’s Airborne Visible/Infrared Imaging Spectrometer-Next Generation (AVIRIS-NG) for six sites spanning the Western Ghats of India. We studied the variation of trait space using principal components analysis at spatial resolutions from the plot level (4m), community level (30m and 100m) to the ecosystem level (1000m). We observed a consistent pattern of trait space across different resolutions, with one axis representing the traditional leaf economic spectrum defined by foliar nitrogen concentration and leaf mass per area (LMA) and another axis representing leaf structure and defense defined by fiber, lignin, and total phenolics. We also observed consistent directionality of environment-trait correlations across resolutions with generally higher predictive capacity of our environment-traits models at coarser resolutions. Among the seven traits, total phenolics, fiber, and lignin showed strong environmental dependencies across sites, while calcium, sugar, and nitrogen were significantly affected by site conditions. Models incorporating site as a fixed effect explained more than 50% of the trait variance at 1000m resolution. LMA showed little dependence on both environment and site conditions, implying other factors such as species composition and perhaps site history strongly affect variation in LMA. Our results show that reliable trait-trait relationships can be identified in coarse resolution imagery, but that local scale trait-trait relationships (resolutions finer than 30m) are not sensitive to broad-scale abiotic/biotc factors.

## Introduction

Foliar functional traits such as specific leaf area and chemical composition play important roles in multiple ecosystem processes related to growth, defense and stress avoidance. They provide an efficient construct for monitoring ecosystem conditions, evaluating biodiversity independent of detailed knowledge of species composition (Jetz et al., 2016; Schneider et al., 2017), and for identifying ecosystem response to a changing climate (Wright et al., 2007; van Bodegom et al., 2014).

Functional trait research has provided insights into tradeoffs between photosynthesis and leaf construction costs, usually referred to as the leaf economics spectrum (Onoda et al., 2017; Wright et al., 2004; Wright et al., 2002, Díaz et al. 2016), as well as into the environmental and evolutionary drivers of plant traits (Díaz et al., 2013; Meng et al., 2015; Wright et al., 2005; Yang et al., 2019). How foliar traits are used to understand vegetation function depends on the scale of analyses, with different interpretations possible from leaf, whole-plant, community, and regional level measurements. Nevertheless, analyses across scales have largely been developed using databases (e.g. TRY, Kattge et al., 2020) built from numerous leaf level studies or local/regional sampling that are neither spatially nor taxonomically comprehensive. Less frequently, studies have explored trait variation and environmental drivers at the community scale using field-collected plot-level data (Bruelheide et al., 2018; Williams et al., 2020). There is now increasing interest in understanding broad scale patterns at the global scale (Boonman et al., 2020; Butler et al., 2017; Migliavacca et al., 2021; Moreno-Martínez et al., 2018; Schiller et al., 2021; Vallicrosa et al., 2022) to better parameterize Earth system models (van Bodegom et al., 2014) and predict impacts of global change. With increasing interest in gridded representations of functional trait variation, there is an increasing need to understand how trait composition changes with scale and what the major environmental drivers of trait variation are at different scales of measurement or aggregation.

A significant challenge faced in community-scale studies analyses is data limitation. Bruelheide et al. (2018), for example, leveraged plot level composition data (over 1.1 M plots), but trait data were imputed using species means from TRY (Kattge et al. 2020) and the authors acknowledged that “using species mean values for traits excludes the possibility of accounting for intraspecific variance, which can also strongly respond to the environment” (p. 1913). Yet, the collection of detailed in-situ data that captures intra-specific and intra-community variation is potentially labor-intensive, expensive, and logistically infeasible for many regions. The quality, representativeness, and comparability of field data also largely depend on sampling protocols, with the potential that different sampling approaches can strongly affect the mean and variance of community-level traits (Asner et al., 2017; Enquist et al., 2017).

Remote sensing approaches-- in particular imaging spectroscopy-- provide an alternative approach to measure some foliar functional traits at regional scales, thus offering the opportunity to study the composition and environmental drivers of traits at multiple scales. Because the data are sparially comprehensive, these approaches open opportunities to better characterize spatial heterogeneity in ecosystem characteristics, although remote sensing generally does not provide species level estimates due to mixed pixels unless the imagery is collected at a fine scale and there is ancillary data enabling identification of species. Early studies were mostly local, and demonstrated the capacity to estimate foliar nitrogen, lignin, and nonstructural carbohydrates from hyperspectral imagery (Martin & Aber, 1997; Matson et al., 1994; Wessman et al., 1989). More recent work demonstrates that data-driven trait models can be applied across different regions and biomes (Martin et al. 2008, Asner et al. 2012, Singh et al. 2015, Wang et al. 2020), and covering a wide range of foliar traits with better than 15% relative error. Existing work covers almost all biomes of Earth, including tropical forests (Asner et al., 2017; Martin et al., 2015), temperate forests (Singh et al. 2015, Wang et al. 2020) and grasslands (Biewer et al., 2009; Mutanga et al., 2004; Ramoelo et al., 2013; Wang et al., 2020) and managed ecosystems (Mahajan et al., 2014, 2017; Moharana and Dutta, 2016; Liu et al., 2021). Studies using imagery-derived trait data have generally confirmed expected trait-environment relationships identified using traditional measurements (Balzotti & Asner, 2018; Martin et al., 2015; Singh et al., 2015). To our knowledge, these studies have not addressed the scale dependency of trait-trait or trait-environment relationships using spatially explicit data.

There are extensive spatial gaps in our knowledge of trait variation, including field measurements from databases (Jetz et al., 2016; Kattge et al. 2020) and from remote sensing data, which to date has largely been provided by airborne imaging spectrometers operated by and in North American and European countries. The Indian subcontinent, for instance, is particularly poorly represented in trait databases (5538 observations in TRY, less than 1% of total observations), despite its large area and diverse habitats. As a consequence, a joint NASA-ISRO campaign to collect Airborne Visible InfraRed Imaging Spectrometer - Next Generation (AVIRIS-NG) imagery over a range of ecosystems in the subcontinent in December-March 2015-2016 and again in 2018 offered the potential to fill an important gap in global knowledge of trait variation. This campaign captured diverse ecosystems along the north-south transect in the sub/tropical Western Ghats mountain belt of India. The Western Ghats stretch from 10°N to 24°N on India’s west coast and covers a large altitudinal gradient from sea level to ca. 2700m. It also covers a broad range of climates with the annual mean temperature ranging from 13.3°C in the high mountains to 28°C at low elevations, and mean annual precipitation varying from ^~^700mm in the rain shadow of the mountains in the east to more than 7000mm along the coast in the west. Diverse environments provide habitat for an extraordinary endemic flora and fauna, which make the Western Ghats one of the eight ‘hottest hotspots’ of biological diversity in the world (Pascal et al. 2004; Rajesh et al. 1996). With high diversities in climate, topography, and vegetation the Western Ghats provide an excellent opportunity to study trait variation, trait-environment relationships, and the scale dependency of these relationships.

Using 4m resolution AVIRIS-NG data, we mapped leaf mass per areas (LMA) and concentrations of foliar nitrogen, sugar, calcium, fiber, lignin, and total phenolics. Among these traits, nitrogen and LMA are correlated with photosynthetic capacity and productivity (Atkin et al., 2015; Madani et al., 2018; Reich, 2014a), with LMA also an indicator of leaf investment in longevity (Poorter, et al., 2009; Westoby et al., 2000; Wright et al., 2002). Calcium is essential to the cell wall and membrane structures, osmoregulation, and signaling (Baribault et al., 2012; Funk & Amatangelo, 2013; White, P. J. & Broadley, 2003). Sugars refer generally to the set of nonstructural carbohydrates (sucrose, glucose, fructose) that are products of photosynthesis; it is used to provide energy for a wide range of metabolic functions and are the building blocks of starches for storage, as well as cellulose and lignin (together, they compose the fiber content) to support cell structures and to defend against herbivores and other physical damage (Weng & Chapple, 2010). Increased foliar sugar content has been identified as an indicator of plant response to water stress, specifically retaining sugars as a reserve of energy rather than investing in building other compounds (He et al., 2020; Liu et al., 2019). Phenolics are secondary metabolites responsible for numerous functions, most notably defense against pests and pathogens (Coley et al., 1985).

We aggregate the trait maps to different scales to identify the scale dependency of both the trait space and the trait-environment relationships. Specifically, we test the following:

1. Are trait-trait relationships and the shape of the trait space conserved during the aggregation from tree/plot level (4m) to community level (30-100m)?
2. At which scales are trait variations better explained by environmental factors--Larger scales (1000m) that represent general environmental patterns or smaller scales (100m and 30m) that characterize the complexity of local terrain?
3. Do trait-environment relationships vary across scales as a consequence of the aggregation of trait values to increasingly coarse resolutions?

## Study site and data collection

### 2.1 Study area

Climatic variability in the Western Ghats leads to a mix of evergreen and dry-deciduous forests. In this study, we collected foliar samples from five sites in the Western Ghats representing different climatic and topographic conditions, as well as from Udaipur north of the Western Ghats (Table 1). Among them, Sholayar experiences high mean annual temperature and precipitation, Mudumalai has a significant temperature gradient caused by its large elevation range (800-2500m), while Muddur is a relatively dry site with moderate temperatures due to its high elevation. To the north, Shimoga shows the greatest precipitation gradient across a narrow temperature range, while the two furthest north sites --Vansda and Shoolpaneshwar—are the hottest due to their low elevations and inland location. Shoolpaneshwar receives much less precipitation than Vansda as it is in the rain shadow of the Western Ghats (Fig. 1).

**Table 1.**
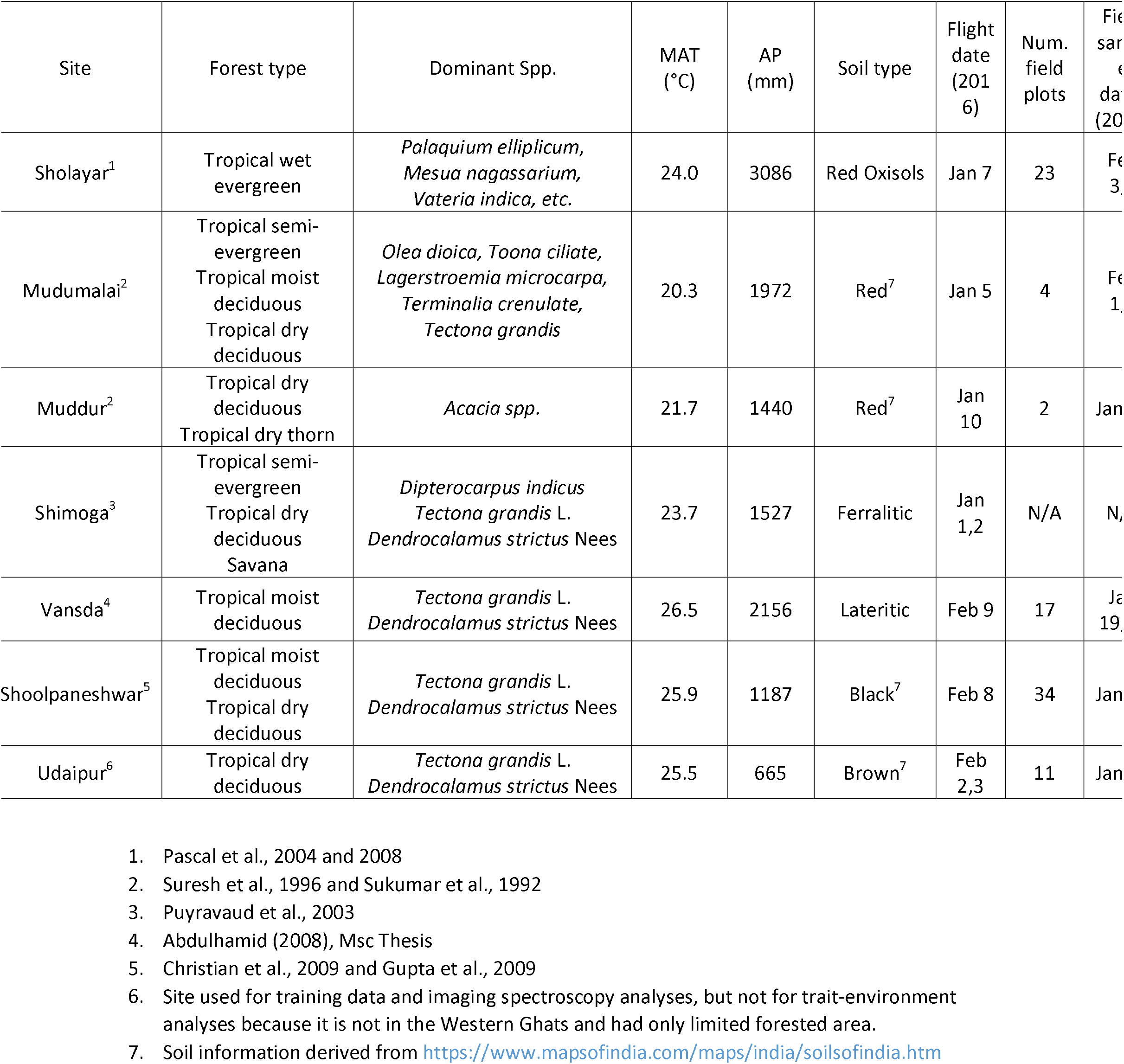
Site characteristics, AVIRIS-NG acquisition times, and field plot information

**Fig. 1.**
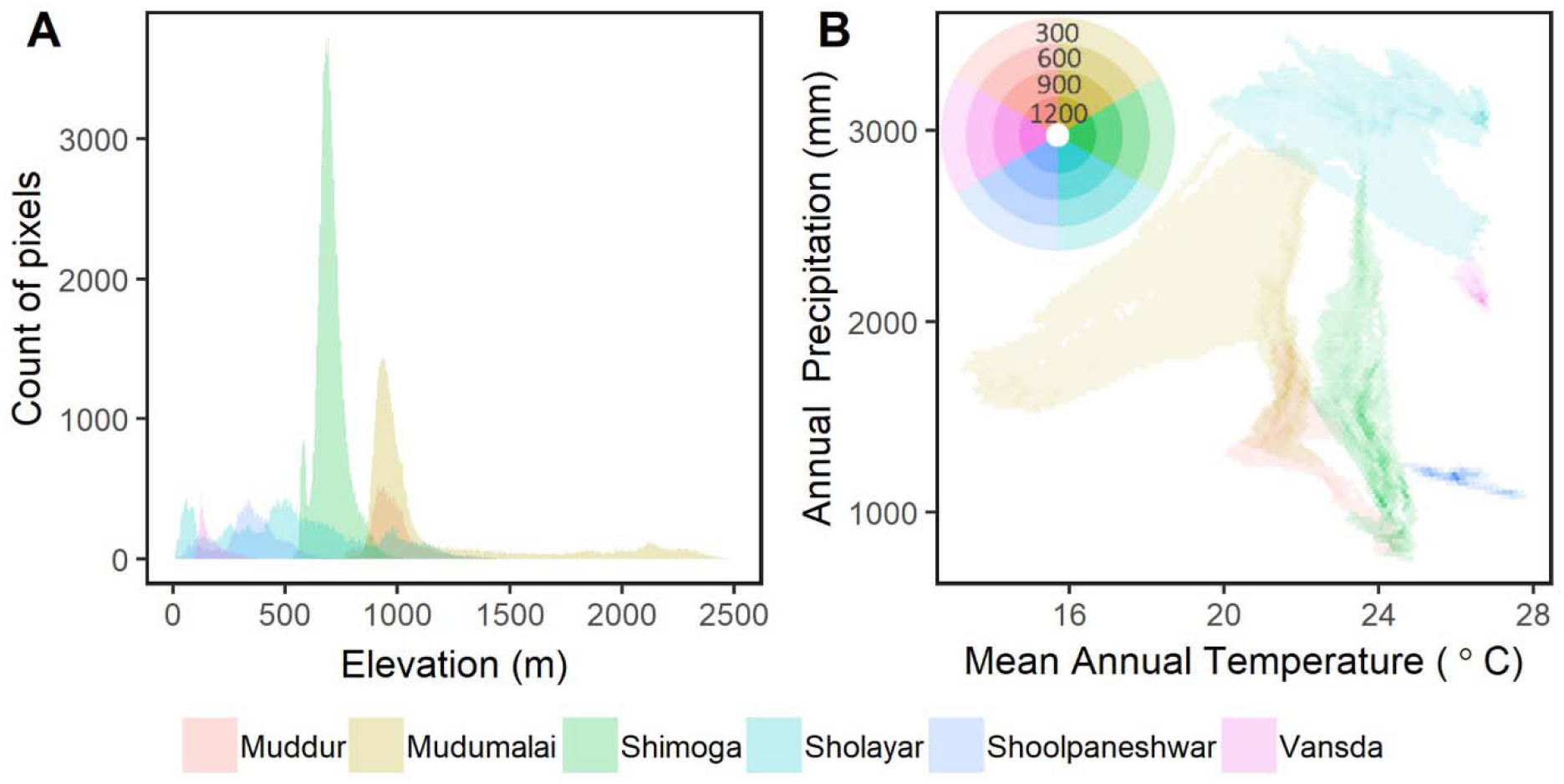
Elevation range (a) and the precipitation-temperature space (b) estimated using data at 100m resolution. Different shades of color in panel **b** represent different number of pixels.

Maps of traits were generated using the models built from field data and imagery from all sites imaged by AVIRIS-NG in the Western Ghats. Data from Udaipur were used for spectral model-building but not included in the environmental analyses due to its limited forest cover.

### 2.2 Foliar sampling and plot trait estimation

Foliar sampling occurred 19 January to 4 February 2018, almost exactly two years following acquisition of AVIRIS-NG imagery in 2016. The use of anniversary foliar samples, rather than concurrent samples, was necessitated for logistical reasons, but was considered reasonable given nearly identical climate records for the two years. Our analyses did not utilize highly labile chemical traits such as pigments or water content, as these may not have been consistent across years. 91 plots from six sites (Fig. 2) were established based on the following criteria: 1) were representative of the local vegetation; 2) were geographically homogenous in a region no smaller than 10m x 10m (i.e., occupying at least four pixels in the 4m resolution AVIRIS-NG image); 3) were accessible with appropriate permissions. Foliar samples were retrieved from the sunlit crown using a combination of pole pruners, slingshot, and in a limited number of plots, manually by trained climbers. We estimated cover percentage and sampled top-canopy foliage for each species in the plot. Latitude, longitude, and elevation of each plot were logged using a portable GPS unit (Garmin GPSMAP 64s) which was later used to locate the site on imagery and delineate sample locations.

**Fig. 2.**
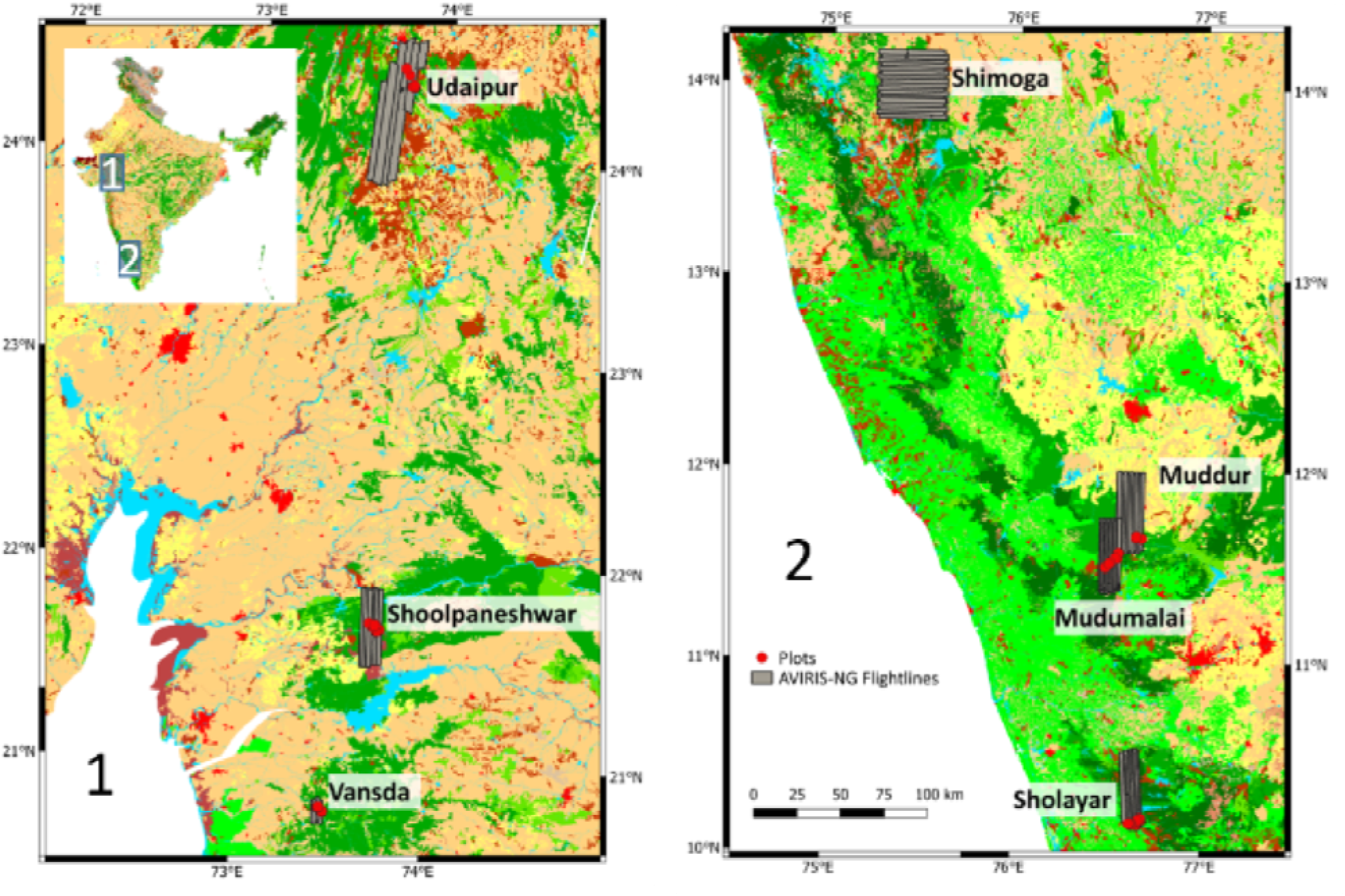
Locations of field plots (91) and AVIRIS-NG flight lines (57) in the Western Ghats of India.

Leaf samples were imaged using a DLSR camera with a reference object of known area (Fig. S1) and left to air dry in a paper bag. Upon return from the field, samples were dried at 70 °C for a minimum of 48 hr and were then weighed to obtain dry mass. Leaf scans were analyzed in ImageJ (https://imagej.net/ImageJ) to obtain leaf area. Leaf mass per area (LMA) was determined by dividing dry mass by projected leaf area. After weighing, the samples were ground to 20 mesh (0.841 mm) using a ball mill and shipped back to the University of Wisconsin-Madison for subsequent determination of foliar biochemical traits.

We measured six spectra of the dry, ground samples with an ASD FieldSpec3 full range (350nm – 2500nm) spectrometer (Analytik, Cambridge, UK) following the protocol from Serbin et al. (2014). Spectra were corrected for ‘jumps’ at 1000nm and 1800nm caused by different detectors used in ASD using the opens source Python package SpecDAL (https://github.com/EnSpec/SpecDAL), and then averaged to yield one spectrum per sample.

We applied generalized partial least square regression (PLSR) models to predict the following foliage traits based on the leaf dry spectra: nitrogen, calcium, sugar, total phenolics, fiber, and lignin. These PLSR models were built using combined data previously reported by Wang et al. (2020) and Serbin et al. (2014). Model performances for the leaf-level traits are summarized in Table S1.

Among the 91 plots, 85 were monospecies in which the top-canopy foliar traits were taken directly as the plot traits. In multispecies plots, we calculated the species-weighted traits based on the cover percentage of each species in the plot.

### 2.3 Hyperspectral image acquisition and processing

In January and February 2016, 57 AVIRIS-NG images from seven sites were acquired along the Western Ghats of India at a flight altitude around 4500m above ground level, yielding a pixel size of around 4m. AVIRIS-NG images 425 bands between 380nm to 2510nm at a 5nm spectral resolution. Data were orthorectified and atmospherically corrected by the Jet Propulsion Laboratory (JPL) following Thompson et al. (2015) and available from https://avirisng.jpl.nasa.gov/dataportal/. The Level-2 reflectance data were then corrected for topographic and BRDF (Bidirectional Reflectance Distribution Function) effects following methods reported by Queally et al. (2021) and openly available at https://github.com/EnSpec/HyTools.

### 2.4 DEM data and topographic gradients

ASTER global elevation data with 30m spatial resolution was downloaded from EARTHDATA (https://earthdata.nasa.gov/) and aggregated to 100m and 1000m, from which slope and aspect were calculated at all three resolutions. We also calculated the topographic wetness index (TWI, Sørensen et al., 2006).

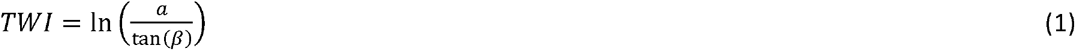

Where *a* stands for the catchment area per unit contour length, and β is the slope. DEM analyses were conducted in QGIS (https://www.qgis.org/en/site/).

### 2.5 Climate and land cover data

Global climate and solar radiation data at 30 arcmin (^~^1km) resolution were obtained from WorldClim Version2 (Fick & Hijmans, 2017; https://worldclim.org), including the average monthly solar radiation, annual mean temperature, average monthly precipitation, and mean annual precipitation for 1970 to 2000. We interpolated the climate data to 100m and 30m resolution based on the corresponding DEM data using radial basis functions as recommended by Ahmed et al. (2014) in the SciPy package of Python 3.7. Based on the monthly precipitation data, we also calculated the seasonality index for precipitation (Walsh & Lawler, 1981) at 30m, 100m, and 1000m resolutions following:

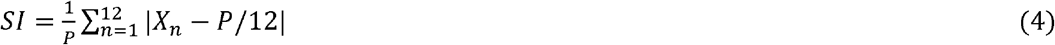

Where P is annual precipitation and X_*n*_ refers to the precipitation of month *n*. Higher values of SI indicate greater seasonality, i.e. precipitation occurring in a few months rather than evenly distributed during the year.

30m resolution land cover data from 2010 for the study area was acquired from GlobeLand30 (http://www.globeland30.org), which is derived from numerous data sources including Landsats 5 and 7 and China’s HJ-1 (J. Chen et al., 2015).

## Data Analysis

### 3.1 Mapping canopy traits using imaging spectroscopy

For each field plot, we extracted spectra from the four AVIRIS-NG pixels nearest to the recorded plot location. We dropped spectra with NDVI < 0.5 (NDVI=(*R*_900_ – *R*_685_)/(*R*_900_ + *R*_685_)) to remove partially vegetated pixels exhibiting substantial background soil influence. In the AVIRIS-NG data, we removed bands in two strong atmospheric water absorption regions (1330 – 1440 nm and 1780 – 1950 nm) and noisy wavelengths below 400 and above 2400 nm. Outlier pixels were removed using principal components analysis (PCA, see S1 in supplementary materials), and remaining pixels were averaged to yield one spectrum per plot. Vector normalization was applied to the spectra to reduce the influence of relative differences in brightness caused by flight direction, atmospheric and sensor conditions across the flight lines (Feilhauer et al., 2010).

Algorithms for mapping foliar traits were developed using permutational partial leastsquares regression to predict field-measured traits as a function of image spectra (PLSR, Wold et.al., 1984). PLSR is widely used where data necessary for radiative transfer models (RTMs) are unavailable, impractical to implement at scale, or suitable RTMs do not exist for particular traits (Asner et al., 2017; Singh et al., 2015; Wang et al., 2019). As a data-driven method, PLSR is effective for handling a large number of input (independent) variables (e.g., 337 bands in this study) with strong collinearity and where the sample size may be comparatively small. It leverages the whole spectrum rather than single bands to estimate chemical traits (Martens, 2001). Here, we trained PLSR models using different ranges of the spectrum—the full range (400-2400 nm), the NIR-SWIR (1000-2400 nm), and SWIR only (1400-2400 nm)—to test whether more restricted ranges yielded more robust models on validation samples by reducing the confounding influence of pigments in the visible and canopy structure in the near infrared region. This is particularly notable for traits like nitrogen, for which the primary spectral features used in estimation are present in the SWIR (i.e., spectral features related to proteins).

We randomly divided the data into calibration (65%) and validation (35%) sets for training and evaluation. We utilized the prediction residual error sum of squares (PRESS) statistic (Chen et al., 2004) to identify the number of orthogonal spectral vectors for PLSR models and prevent overfitting. We permuted the analyses to generate 500 PLSR models for each trait following Singh et al. (2015) and used the following metrics to evaluate performance: The coefficient of determination (R^2^), the root mean squared error (RMSE), the normalized RMSE (NRMSE = RMSE/trait range), and the bias.

PLSR coefficients from the 500 models were then applied to the AVIRIS-NG images to map traits with the mean prediction per pixel as the trait estimate and the standard deviation (SD) as a pixel-wise measure of uncertainty. Due to the strong dependence of the SD on the absolute values of different traits, it is difficult to compare the uncertainties across traits. Thus, we used relative uncertainties (standard deviation/mean) following Verrelst et al. (2016) to screen uncertain pixels in subsequent analyses, using a 25% uncertainty threshold. As well, we filtered out: 1) shadowed pixels, based on their brightness level across the near-infrared region (1017 nm – 1097nm, see S2 for details); and 2) pixels with low vegetative cover or high soil fraction, using NDVI < 0.5. Finally, we used GlobeLand30 to mask non-forest pixels.

We aggregated the cleaned trait maps to 30m, 100m, and 1000m resolutions for each site for further analysis.

### 3.2 Relationships among traits in the study area

We used PCA and Pearson correlation to assess the relationships among traits at the leaf level, plot level (using native ^~^4m AVIRIS-NG trait maps), community level (aggregated to 30m and 100m), and the ecosystem level (1000m). The 30m and 1000m grid cells may be of particular interest because they represent resolutions that may be expected for forthcoming hyperspectral satellite missions such as SBG (30m) and a resolution commonly used for terrestrial ecosystem models (1000m). For the 4m data, we randomly extracted ^~^1000 pixels per flightline for analysis, resulting in a dataset for statistical analyses of 7980 valid pixels after masking. All the valid pixels were used in the analysis at coarser resolutions.

### 3.3 Environmental drivers of trait variation at different scales

We considered six environmental predictors of trait variation that cover hypothesized heat, light, water, and topographic drivers: mean annual temperature (MAT), annual solar radiation (radiation), mean annual precipitation (MAP), precipitation seasonality (SI), elevation, and TWI. We then calculated the variance inflation factor (VIF) to test the multicollinearity among the predictors and removed elevation due to its high correlation with MAT (Table S2). The analyses were based on 2000 randomly selected valid pixels per site at 30m, 500 at 100m, and 30 at 1000m, with sample sizes being a consequence of study area extents and the forest coverage in the study area.

We built linear models (LMs) to explain trait variation using environmental predictors. During this process, an autocovariate term was introduced into the linear models to account for the spatial autocorrelation in the selected samples. We calculated the autocovariate using the ‘spdep’ package in R following Bardos et al. (2015).

Among all the traits, total phenolics followed a log-normal distribution and was natural log transformed. The environmental predictors were zero-centered to reduce multicollinearity (Du et al., 2020).

With the linear models, we further evaluated the relative importance of each predictor by its contribution to the model R^2^. Due to the existence of collinearity among our predictors, the contribution of R^2^ from each predictor will be affected by its order in entering the equation if estimated using the stepwise regression. We thus partitioned R^2^ using the average stepwise method (Lindeman et al., 1980) which permutates all possible entering orders and calculates the average contribution of R^2^. The resulted R^2^ partition represents the predictive power for each predictor (Bring, 1995). This analysis was carried out using relaimpo package (Grömping, 2006) in R. To further isolate the influence of each predictor on the trait variation, we applied partial regression to LMs (car package in R, Fox & Weisberg, 2019). Finally, we incorporated ‘site’ as a fixed effect into the LMs to explore the site effects on the trait variation.

## Results

### 4.1 Trait models

PLSR models for the seven traits produced calibration R^2^ ranging from 0.54 to 0.9 and NRMSE between 8% to 17% (Table 2, Figure 3), with the best calibration performance for sugars content (R^2^ = 0.9 and NRMSE of 8%), followed by calcium, LMA, and fiber content with ^~^0.7 R^2^ and NRMSE within 15%. Calibration results for total phenolics, nitrogen content, and lignin were moderate, with R^2^ > 0.5 and NRMSE < 20%, indicating that these models characterized the variation of these traits but with lower precision than the others. Sugar content also exhibited the best validation performance (R^2^ = 0.67, NRMSE = 15%), with other traits validating with R^2^ around 0.5 and NRMSE about 20%. The poorest performance was from fiber content (R^2^ = 0.43, NRMSE = 21%). For most traits, PLSR models using the whole spectrum yielded the best performance, although models utilizing the 1000-2400 nm NIR-SWIR bands performed best for nitrogen, fiber, and lignin (Table 2, see Table S3 for all comparisons), which was unsurprising given that most known absorption features for nitrogen (e.g., 1510, 2060, 2180 nm), fiber (1120, 1200, 1780, 2270 nm), and lignin (1120, 1420, 1690, 2102 nm) are situated in the NIR-SWIR region (Curran, 1989; Kokaly et al., 2009; van der Meer, 2004). Thus models using this region capture essential information while reducing irrelevant or confounding signals in the visible and red-edge regions. Overall, our PLSR model performance was comparable to results reported for similar traits by Asner et al. (2015), Singh et al. (2015), and Wang et al. (2019, 2020).

**Table 2.**
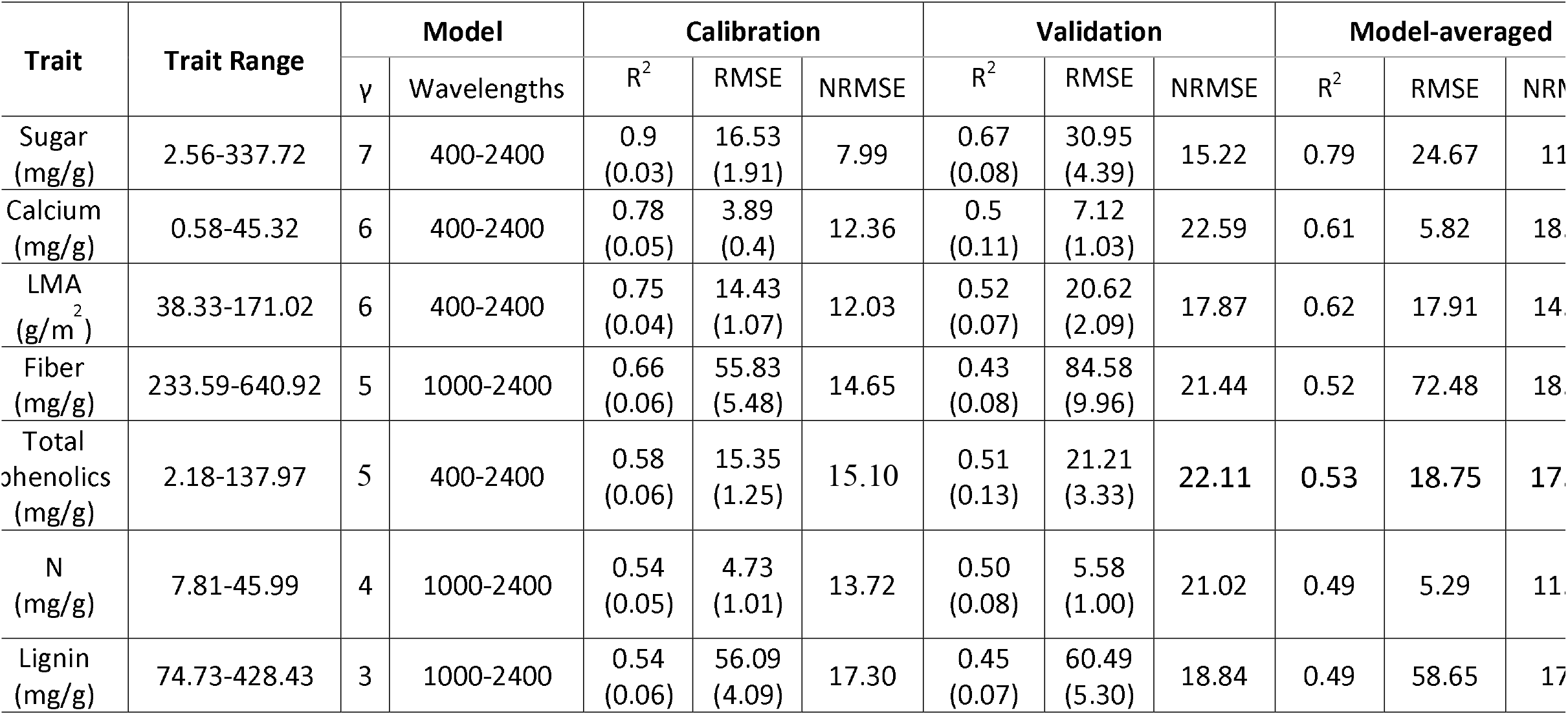
Trait ranges for field samples and summarized performances for PLSR (partial least squares regression) models, γ refers to the number of components in the model.

**Fig. 3.**
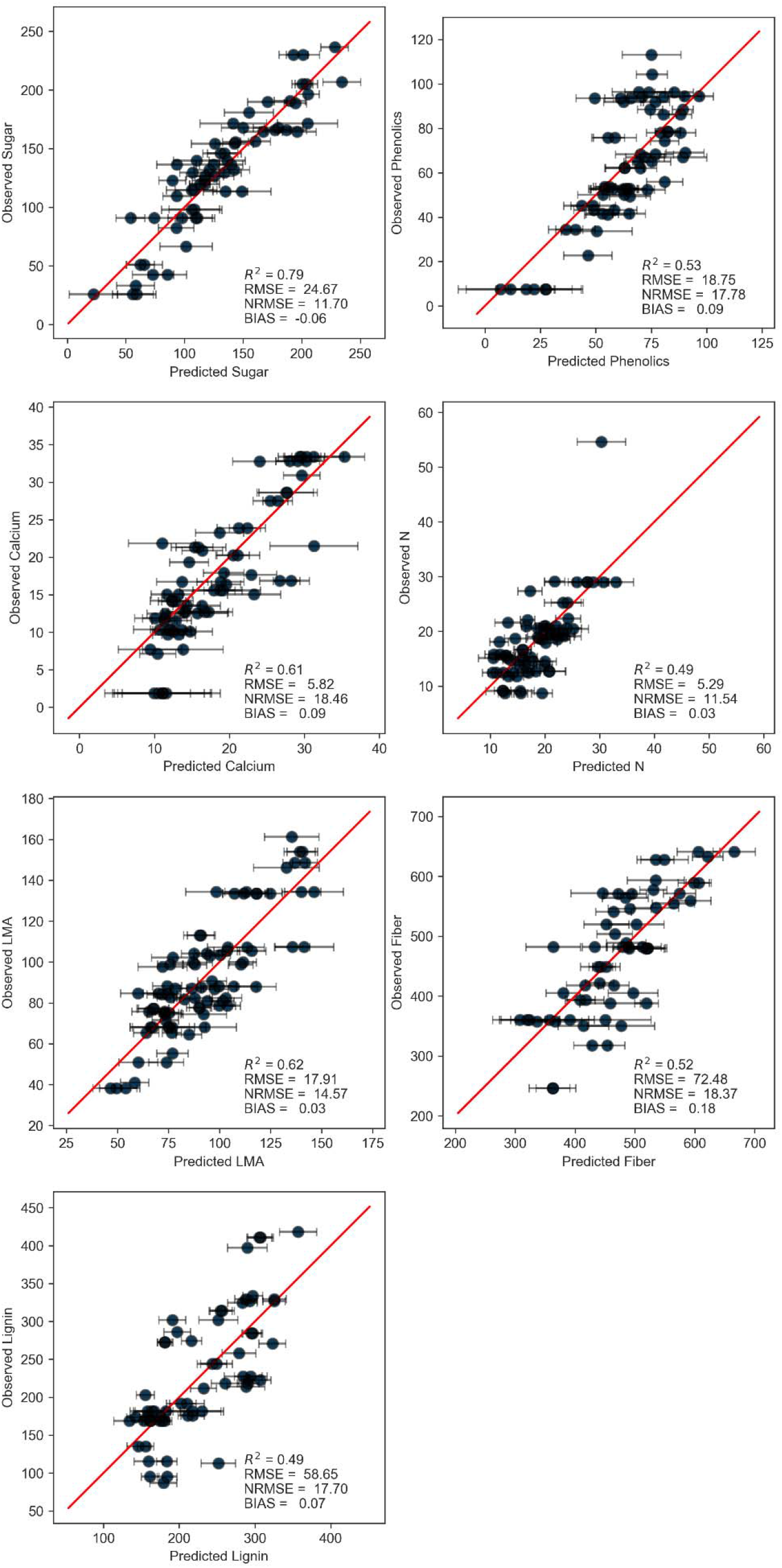
PLSR model fits between the AVIRIS-NG predicted traits and the field measured plot traits. Horizontal error bars indicate the uncertainty (1 SD) of the prediction based on 500 models. Statistics on the plots show the model performances when applied to the whole dataset (Model-averaged in Table 2).

### 4.2 Patterns of trait distributions

The range of trait values across each site was similar, except for lignin (Fig. 4). For most traits, the site mean was within 13% of the regional mean for the Western Ghats study area except for lignin of Sholayar and Shimoga, sugars and calcium of Shoolpaneshwar, and foliar nitrogen at Mudumalai.

**Fig. 4.**
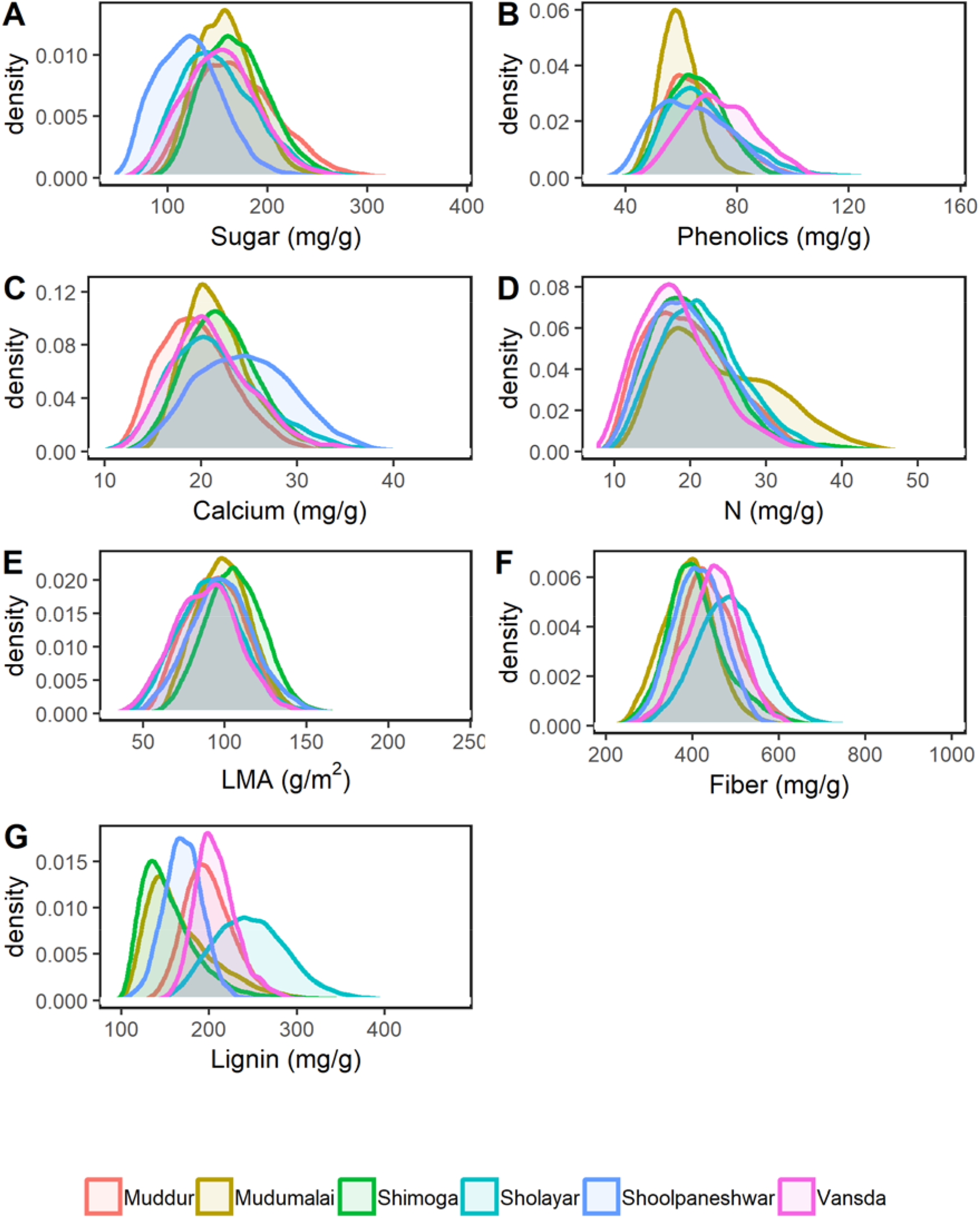
Distributions of traits estimated based on randomly extracted pixels (n = 140 per flight line) for each site.

The site mean of lignin was 30% larger than the regional mean for Sholayar, the wettest sites in the study area (foliar lignin of 250 mg g^−1^ for Sholayar, compared to 190 mg g^−1^ for the whole region). The pattern was also apparent on a PCA plot of traits for the region (Fig. 5), where Sholayar delineated an axis representing lignin (cyan circle in Fig. 5). Shimoga, a comparatively dry site, had 17% lower lignin than the regional mean.

**Fig. 5.**
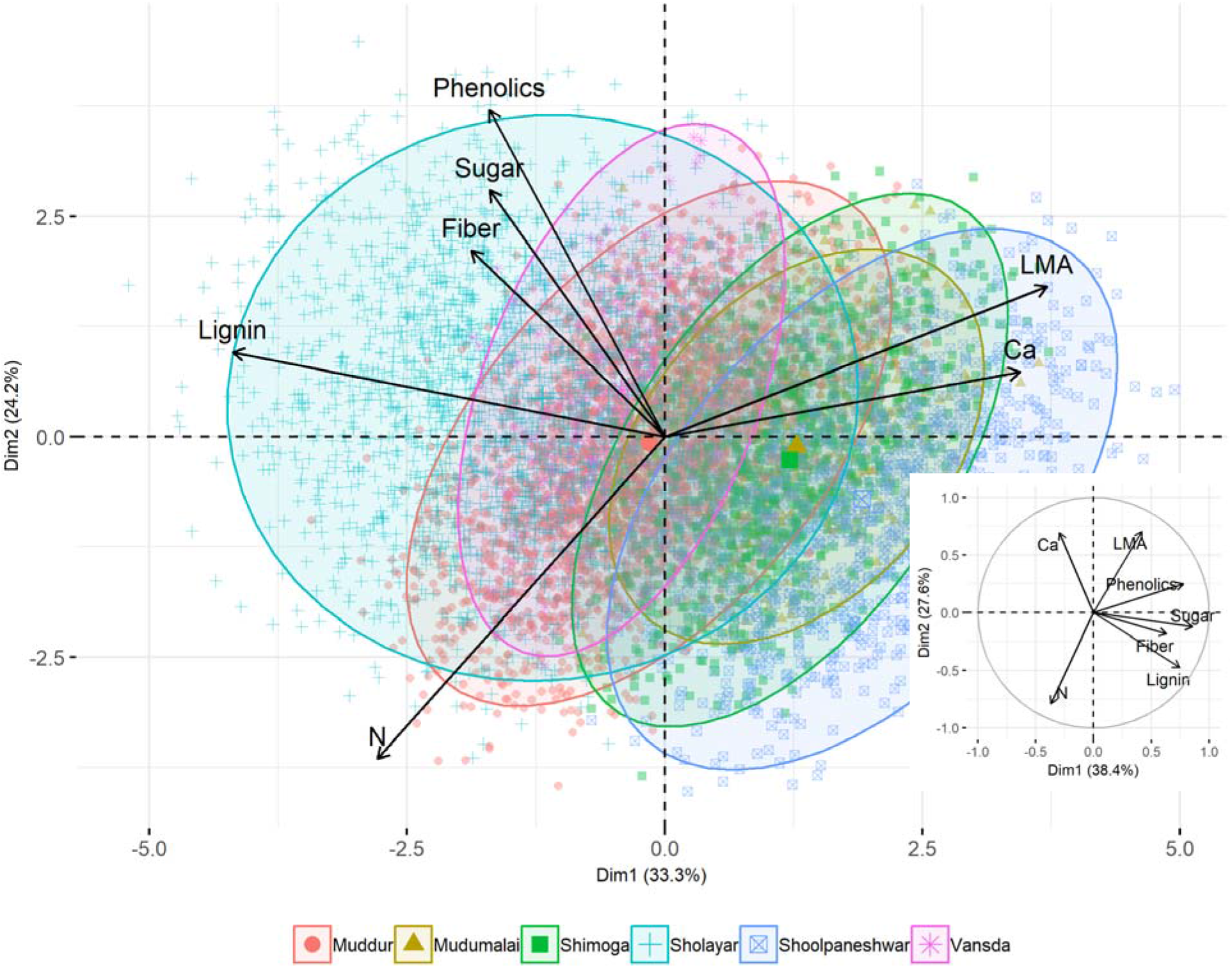
Principal component plots for image-level traits (^~^4m resolution) and the leaf-level traits (inset). Colored ellipses outline the trait space for different sites.

Shoolpaneshwar, the driest site retained in the environmental analyses, exhibited 24% lower sugar concentration and 13% higher calcium than the regional mean, while Mudumalai had 14% higher nitrogen concentration.

Most traits exhibited a unimodal distribution (Fig. 4), except for foliar nitrogen at Mudumalai, which was double peaked at 20 mg g^−1^ (roughly equivalent with the other sites) and 30 mg g^−1^. Further investigation revealed large areas of tea plantations within the Mudumalai flight boxes, accounting for the areas that contributed to the second peak in the density plot having higher foliar nitrogen concentration.

### 4.3 Pair-wise relationships among traits

Pearson correlations at the leaf level (Fig. 6A) revealed strong negative correlations between N and LMA (r = −0.51, *p*< 0.01), consistent with the general tradeoff between acquisitive and conservative strategies that are the basis of the leaf economic spectrum concept (Díaz et al., 2016; Onoda et al., 2017; Reich, 2014b; Wright et al., 2005, 2004). Lignin showed a strong positive relationship with fiber, as it is a constituent of fiber assays (r = 0.72, *p* < 0.01). The photosynthetic product sugar was positively correlated with total phenolics, generally considered to be defensive compounds (r = 0.66, *p* < 0.01) as well as the structural compound lignin (r = 0.61, *p* < 0.01), which in combination can be interpreted as investments in leaf longevity and defense.

**Fig. 6.**
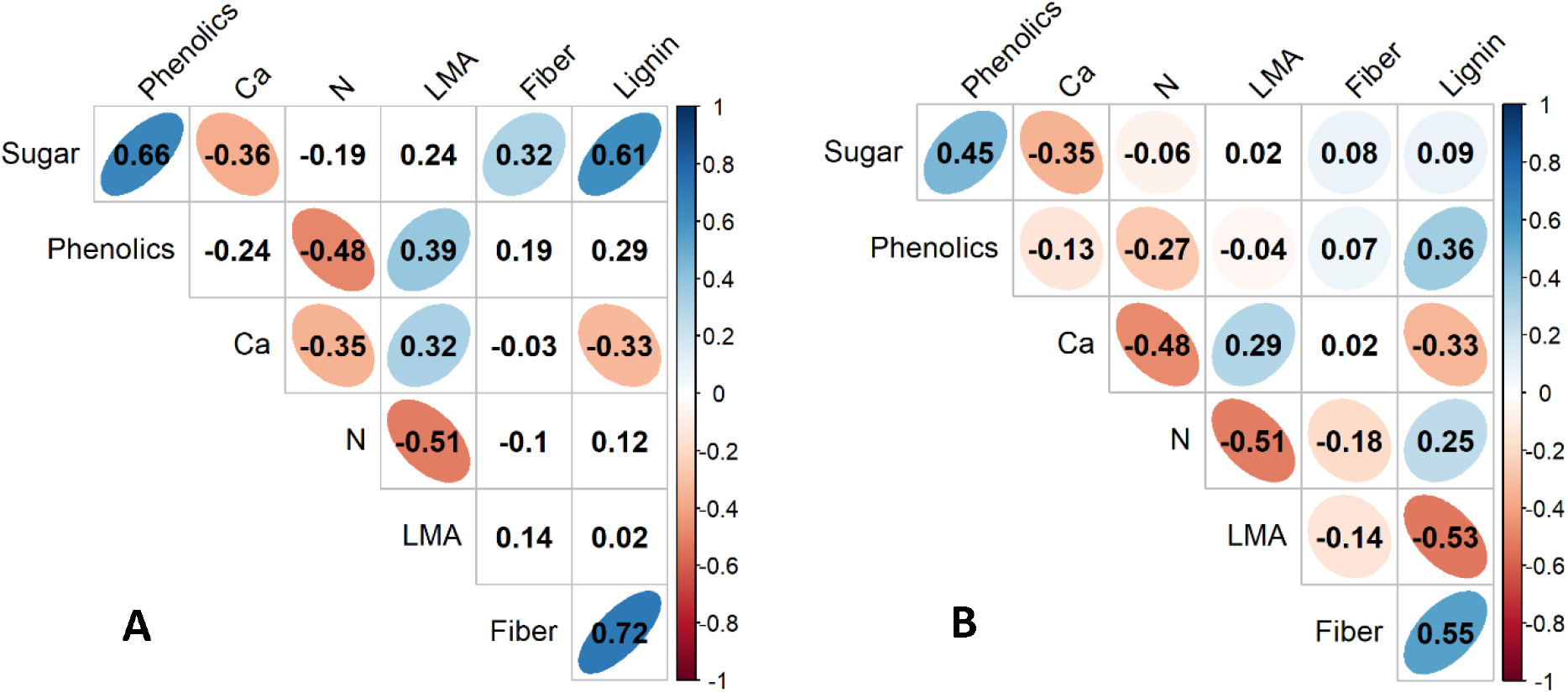
Pearson correlation coefficients between traits for **(a)** leaf level and **(b)** image predicted traits at ^~^4m resolution. Ellipses are drawn around correlations with *p*<0.01.

The correlations among image-level traits at ^~^4m resolution (Fig. 6B) displayed similar patterns as the leaf-level traits. Some weak relationships observed at the leaf level became stronger, such as the positive correlations between total phenolics and lignin. In contrast, the strong positive correlation between sugar and lignin (also fiber) diminished at the image level. Those differences can be attributed in part to sample size (larger number of samples from image data); also, image pixels often comprised multiple species and represented an integrated signal of foliar properties.

The first two components of a principal components analysis explained 66% of the leaf-level variation and 57.5% at the image level (Fig. 5). Two orthogonal axes for image-level traits represent one gradient of high N to high LMA and the second axis of increasing fiber, sugar, total phenolics, and lignin. The first axis is closely related to photosynthesis and leaf morphology and thus the leaf economics spectrum (Wright et al., 2004), while the second axis represents structure-defense traits. The third component from our PCA analyses (both leaf and image level) further groups fiber, lignin, and calcium on one end and sugars and phenolics on another (the vertical axis on Fig. S2). Image level traits preserved the patterns we observed from the leaf level traits (the inset on Fig. 5), confirming the general concurrence of the trait maps with field samples.

Correlations and principal components analyses at 30m, 100m, and 1000m (Figs. S3 and S4) yielded similar results as the native resolution, indicating the relationships among traits were well maintained when data were aggregated at scales from the single plots (4m) to the ecosystem scale (1000m). Notably, the first two components were able to explain ^~^70% of the total trait variation at 30m, 100m, and 1000m, almost 15% more than at the leaf- and plot-level. Such results revealed strong convergence in trait variance when aggregated from the plot level (^~^4m) to the community level (30m), and illustrate a level of stochasticity that may occur at fine scales.

### 4.4 Trait-environment relationships

#### 4.4.1 Modeling trait variation using linear models

Our study sites along the Western Ghats cover a wide range of elevation and climate (Fig. 1). The linear models (LMs) predicting traits as a function of environmental predictors performed the best at the coarsest (1000m) scale, except for foliar sugar concentration (Table 3 and Fig. 7). Total phenolics and lignin were well predicted at all three resolutions (R^2^ 0.37 – 0.66). The LMs for fiber were moderate at 30m and 100m, but explained 51% of the variance at 1000m. Models for calcium showed moderate performances for all resolutions (R^2^ 0.21-0.30), with one-third of the variation explained at 1000m. The worst models were for nitrogen and LMA, where environmental predictors only explained 15-25% of the variation across all scales.

**Table 3.** Model R^2^ for linear models at different resolutions.

| | Sugar<br>(mg/g) | ln(Phenolics)<br>(mg/g) | Calcium<br>(mg/g) | N<br>(mg/g) | LMA<br>($\text{g/m}^2$) | Fiber<br>(mg/g) | Lignin<br>(mg/g) |
| --- | --- | --- | --- | --- | --- | --- | --- |
| 30m | 0.21 | 0.37 | 0.21 | 0.14 | 0.15 | 0.29 | 0.50 |
| 100m | 0.27 | 0.41 | 0.24 | 0.18 | 0.16 | 0.35 | 0.55 |
| 1000m | 0.12 | 0.66 | 0.30 | 0.25 | 0.20 | 0.51 | 0.59 |

**Fig. 7.**
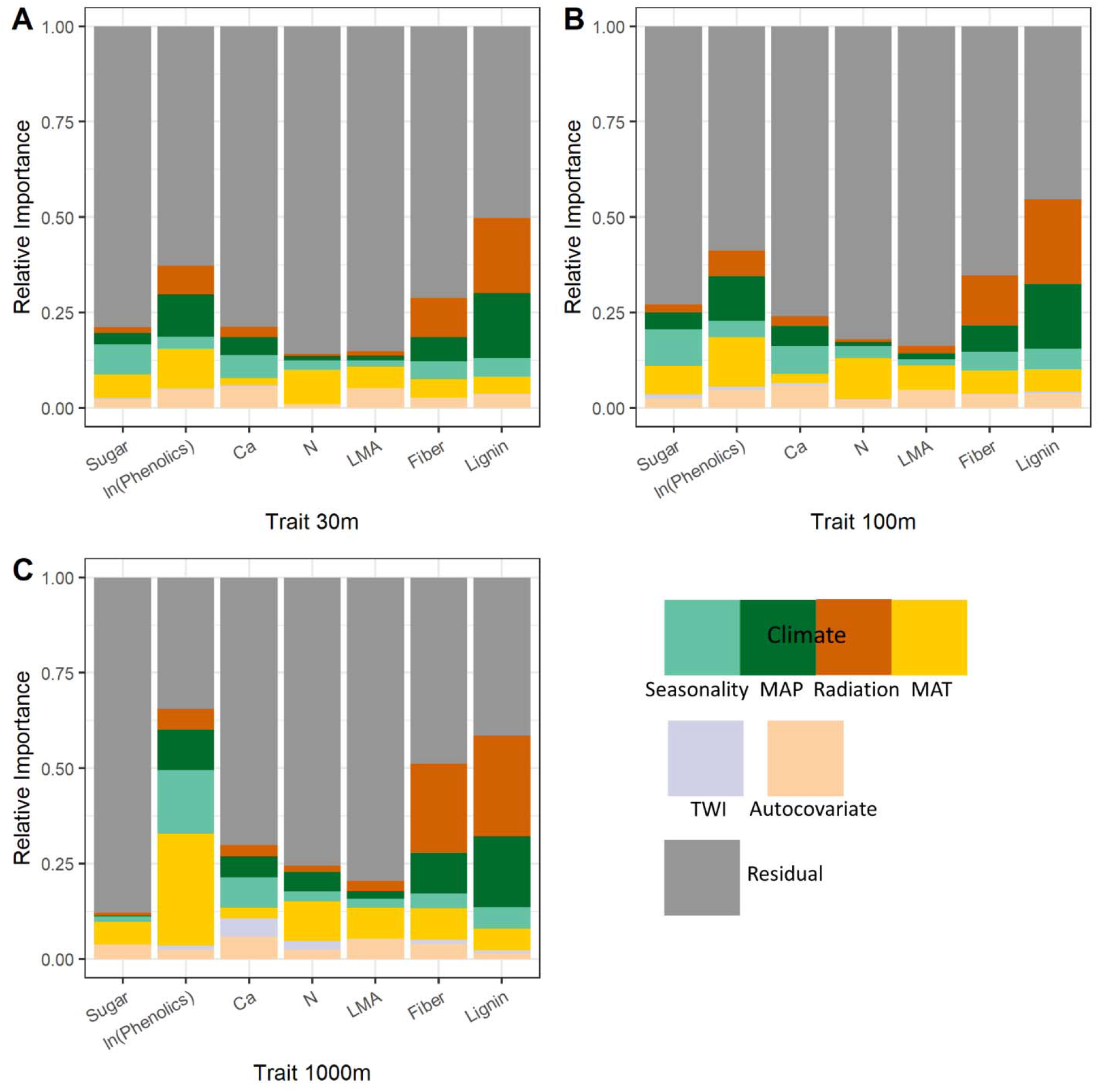
Relative importance for environmental predictors at three resolutions.

The relative contribution of each predictor to model R^2^ is listed in Table S4. For the best-predicted traits—phenolics, fiber, and lignin, MAT contributed the most to the prediction of phenolics models at 100m and 1000m (32% and 45% of R^2^). In contrast, solar radiation appeared to be the most important predictor for fiber and lignin, contributing more than 35% to the model R^2^. Seasonality was the strongest predictor for foliar calcium (all resolutions) and sugars (30m and 100m). MAT contributed the most among environmental variables in models for nitrogen. Among all the traits, LMA and calcium exhibited the strongest spatial correlation with the autocovariate contributing to more than 20% of the models’ R^2^. TWI was the weakest predictor in the models, only accounted for less than 5% of the R^2^ in most cases. For a given trait, the relative contributions of a predictor were generally stable across different resolutions.

#### 4.4.2 Partial regression analyses

We applied partial regression to further isolate the unique relationship between a given trait and an environmental variable while controlling for other predictors (Fig. 8). Total phenolics increased significantly with higher MAP, hotter MAT, and stronger solar radiation (*p*<0.01, all resolutions), but decreased with precipitation seasonality (*p*<0.01, 30m and 100m). Fiber and lignin varied similarly with most environmental variables, decreasing with stronger solar radiation and greater precipitation seasonality (*p*<0.01, all resolutions) while increasing with higher MAT (*p*<0.01, all resolutions). Mean annual precipitation exhibited a moderate positive relationship with lignin (*p*<0.01, 30m and 100m) but correlated negatively with fiber (*p*<0.01, 100m). In general, lignin and fiber were higher in hotter places with lower solar radiation and weaker seasonality. This is most notable in the higher lignin and fiber concentration in the foliage from Sholayar (Fig. 4 and Fig. 5), the furthest south and wettest site.

**Fig. 8.**
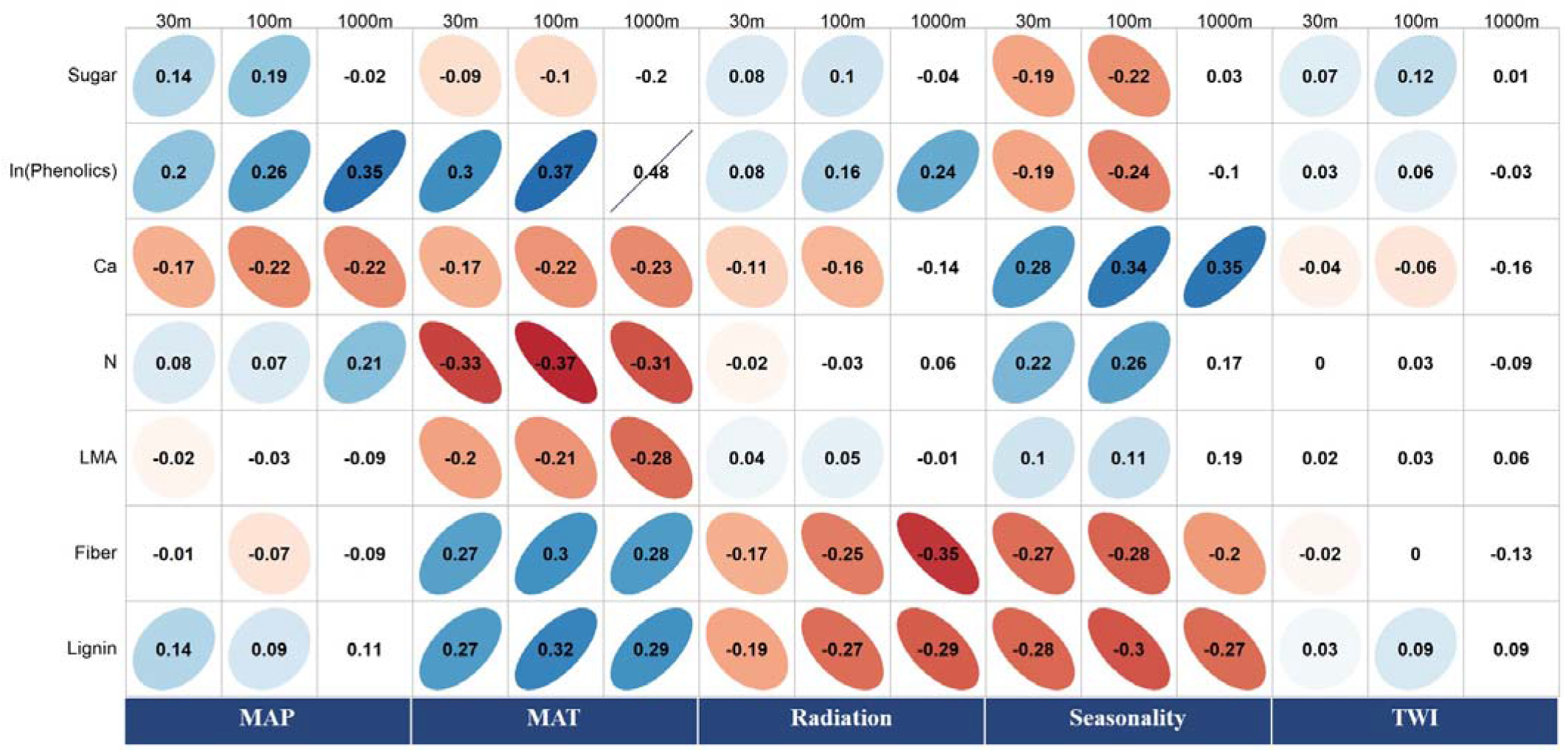
Partial correlation plots between traits and environmental variables at different resolutions. Ellipses are drawn around correlations with *p*<0.01.

Although environmental variables explained less than 30% of the trait variation for calcium, LMA, nitrogen, and sugars, partial regression analysis provided useful insights that support findings from previous studies. For example, foliar nitrogen was positively correlated with MAP, which has been observed in both remote sensing and traditional analyses (Asner & Martin, 2016; Körner et al., 1986; Meng et al., 2015). A negative influence of MAT on leaf nitrogen concentration was also observed, which may link to changes in species composition along the altitudinal gradients in which species with higher foliar nitrogen are found at higher elevations with lower MAT (Balzotti & Asner, 2018; Morecroft et al., 1992), thus representing a change in traits due to species turnover rather than within-species variation (Dong et al., 2020). The identification of positive influences of MAP and solar radiation and negative influence of seasonality (at 30m and 100m resolutions) on foliar sugars is a new insight, although these relationships may also be tracking the correlation between sugars and phenolics. Calcium was positively associated with precipitation seasonality (*p*<0.01, all resolutions) but negatively correlated with MAP, MAT (*p*<0.01, all resolutions), and solar radiation (*p*<0.01, 30m and 100m), which may be a consequence of Ca availability since weathering would be expected to be higher with greater insolation, higher precipitation, and hotter temperature.

#### 4.4.3 Site effects on the trait variation

Inclusion of site as a fixed effect in the linear models (denoted as LM-site) improved model predictions (Table 4). Most significant improvements were found for sugars, calcium, and nitrogen, where LM-site was able to explain at least 49% of the trait variation at 1000m. Site alone contributed at least 40% of the model R^2^ at all resolutions. Moderate improvements were observed for LMA, where LM-site managed to explain 22%-31% of the trait variation comparing to 15%-20% by the original LMs. Statistical models for phenolics, fiber, and lignin benefitted the least from including site. In most cases, site contributed to less than 35% of the R^2^, especially for lignin (6%) and phenolics (9%) at 1000m.

**Table 4.** Model performance (R^2^) after including site as a fixed factor. Percentages in parentheses indicate the improvements in R^2^ brought by site.

|  | Sugar<br>(mg/g) | ln(phenolics)<br>(mg/g) | Calcium<br>(mg/g) | N<br>(mg/g) | LMA<br>(g/m<sup>2</sup>) | Fiber<br>(mg/g) | Lignin<br>(mg/g) |
| --- | --- | --- | --- | --- | --- | --- | --- |
| 30m | 0.51(59%) | 0.55(33%) | 0.48(56%) | 0.27(47%) | 0.22(31%) | 0.38(24%) | 0.57(12%) |
| 100m | 0.62(57%) | 0.63(35%) | 0.58(59%) | 0.30(40%) | 0.23(32%) | 0.47(26%) | 0.61(10%) |
| 1000m | 0.51(77%) | 0.72(9%) | 0.68(56%) | 0.49(49%) | 0.31(35%) | 0.67(24%) | 0.63(6%) |

## Discussion

### 5.1 Change in variance and conservation of trait space during aggregation

The primary objective of our paper was to evaluate trait-trait and trait-environment relationships across scales in the Western Ghats of India, given accurate maps.

We observed a contraction in trait variance through the aggregation for all traits across all sites (Fig. S5 and Table S5), although trait space was generally conserved. With all sites pooled together, for most traits, trait variance decreased most during the aggregation from plot/tree-level (4 m) to community level (30 m), followed by the aggregation from 30 m to 100 m. From 100 m to 1000 m, trait variance declined much less (Fig. 9). The reduction in trait variance during aggregation differed among traits and among sites for a given trait, likely caused by site-specific differences in patterns of spatial distribution in traits. Unexplained site-specific differences may be a consequence of unmeasured variation related to geology or land use history. The nitrogen maps for Mudumalai and Sholayar are good examples of such effects on trait variance (Fig. 10). Mudumalai was characterized by a low foliar nitrogen region in the north and high nitrogen in the south. As a result, nitrogen variance in Mudumalai was well preserved during aggregation—the 1000m data still contained almost 55% of the original variance at 4m. In contrast, Sholayar displayed a patchy pattern with alternating high and low nitrogen areas. During the aggregation, hotspots of both high and low areas were blended, causing a rapid contraction in the variance of nitrogen in which only 23% of the original variance at 4m was present at 1000m. Areas with heterogeneous (patchy) trait distribution suffer most from information loss during aggregation. Depending on applications, trait maps at coarse resolutions should be used with caution for such areas. For use in applications such as terrestrial ecosystem models, trait maps at resolutions such as 1000m should incorporate information on trait variability as well as trait averages.

**Fig. 9.**
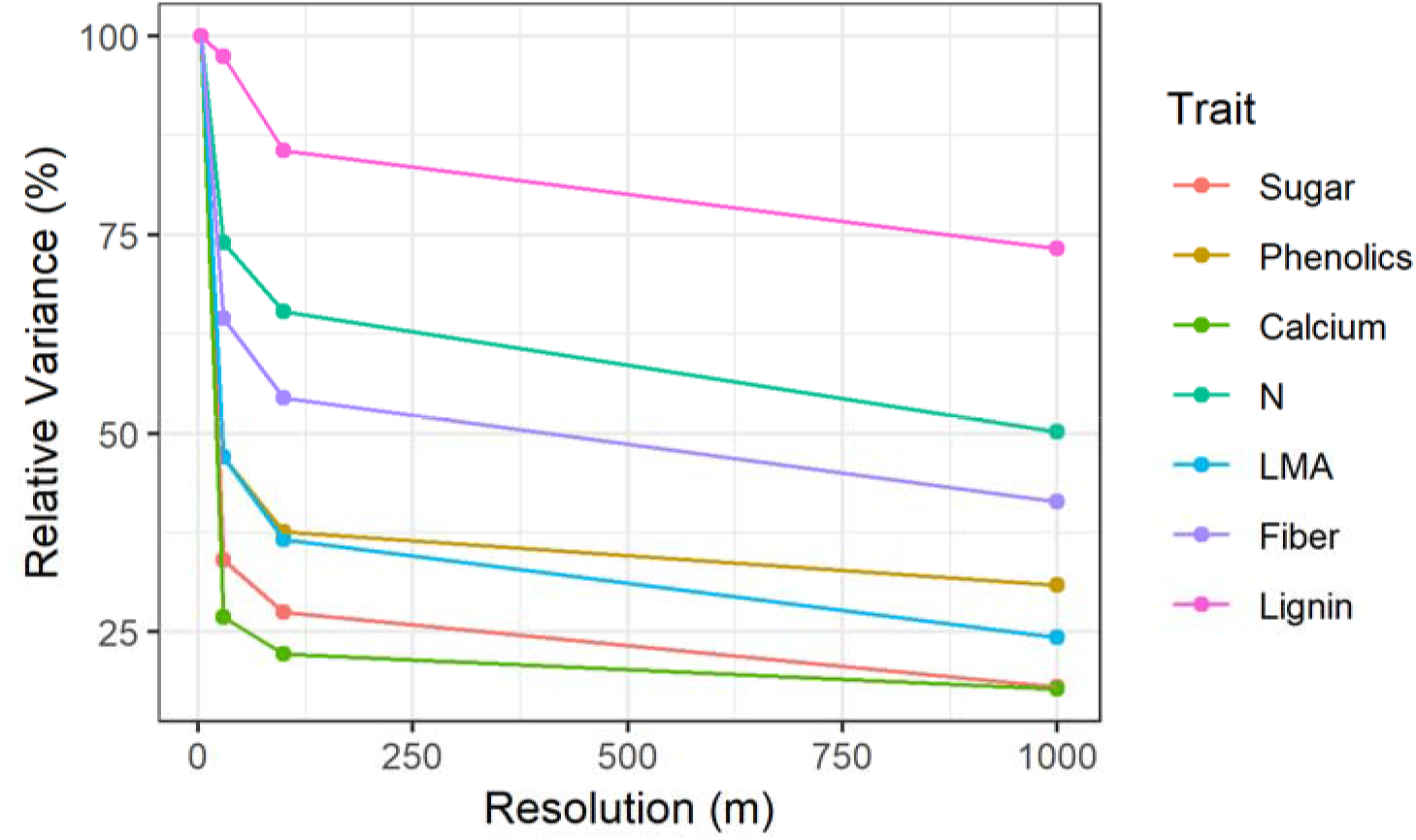
Trait variance at different resolutions. Shown as the percentage of trait variance at 4m resolution.

**Fig. 10.**
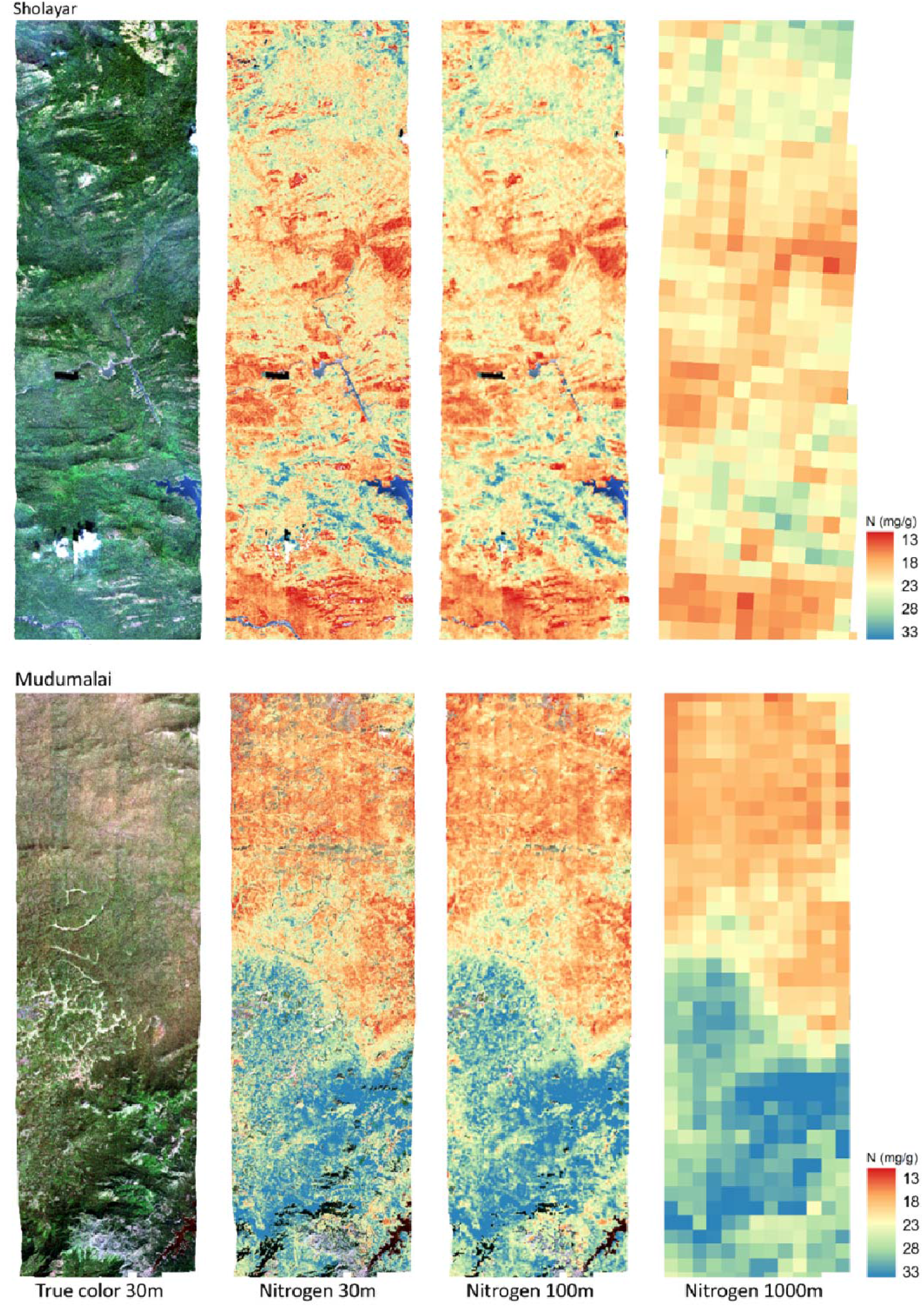
Aggregation effects on trait variance using nitrogen maps for Sholayar (top) and Mudumalai (bottom) as examples. The same color scheme was applied to all the trait maps to improve comparability.

Despite the uneven changes in variance deduction among traits at different sites and across resolutions, the overall trait space was well preserved when all data were pooled. First, the pair-wise correlation between traits exhibited a high degree of agreement across resolutions, especially from 30m to 1000m (Fig. S3). Second, our PCA analyses yielded similar trait space at different grain sizes (Fig. S4). Our findings echoed the observation from Bruelheide et al. (2018) based on 1.1 million global plots where similar tradeoffs between traits were found at both species and community levels.

### 5.2 Trait-environment relationships across resolutions

Our models revealed that environmental variables explained more variation in traits at the coarsest resolution of aggregation (1000m), suggesting that aggregation homogenizes local variation and emphasizes the broader patterns represented by our coarse-scale environmental predictors (Table 3).

The direction and relative strength of influence of almost all environmental variables were preserved during the aggregation of traits. Specifically, the models remained stable at different resolutions with similar contributions to model R^2^ at all resolutions from the same predictors (Table S4). As well, the partial regression analysis (Fig. 8) revealed similar correlations between a given trait and environmental variables across scales.

Environmental variables exhibited the strongest control on fiber, lignin, and total phenolics among the seven traits we studied. These traits are important to plant defense, with lignin (especially) and fiber being important structural compounds that are unpalatable to herbivores and thus act as physical defensive substances. Assays for total phenolics measure antioxidant capacity and comprise numerous compounds that perform important roles in chemical defense. Higher concentrations of fiber, lignin, and total phenolics were associated with hotter environments with lower precipitation seasonality, possibly in response to greater herbivore pressure in these environments (Hallam and Read, 2006). Annual solar radiation influenced fiber and lignin differently than total phenolics: leaf fiber and lignin concentrations decreased with increasing solar radiation. This trend may be related to the resource availability hypothesis (Coley et al., 1985), namely that in places with high solar radiation with all other conditions being equal, plants invest more in photosynthetic machinery rather than defense to achieve faster growth. This topic requires further study, as there are few studies on the effects of solar radiation on foliar fiber and lignin at broad spatial scales or using remote sensing derived estimates. Increases in phenolics with solar radiation can be attributed to a defensive response—intense solar radiation induces ultraviolet stress and triggers the production of phenolics (Wang et al., 2016).

Including site as a fixed effect in the LMs yielded the most significant improvements for predictions of foliar calcium, nitrogen, and the photosynthetic product—leaf sugar concentration--suggesting that site-specific factors strongly control these traits in the Western Ghats. First, site likely captures climate variations that are not expressed in our broad-scale climate data. Site level variation may affect acclimatization and competition among species (Asner & Martin, 2016), leading to differences in nutrient uptake, photosynthesis, and the allocation of its products. Second, site effects also are a surrogate for differences in geology and soils that influence nutrient availability (such as Ca, e.g., Asner et al., 2014) and nitrogen cycling. Among our study sites, Shoolpaneshwar is located on top of black soil rich in calcium carbonate, while other sites share a similar red soil bed containing little calcium due to strong weathering (Table 1). As a result, we observed higher foliar calcium in the AVIRIS-NG derived trait maps from Shoolpaneshwar (Figure 4C). Finally, site effects may capture differences in land use history and species composition (or phylogeny) that affect traits but are not otherwise represented in the data.

LMA was not well predicted as a function of environmental variables; the 1000m scale model (with site effect included) explained only 31% of the variation in LMA. Our LMA dataset exhibited comparable variation (ranges: 33.7-240.6 g m^−2^) to other datasets reported from the tropics (e.g., Asner et al. 2011, Serbin et al. 2019), so this result does not arise from a lack of variation. LMA variation was comparable across sites (Fig. 3E), but little variation was explained by the environment, indicating that species with comparatively low and high LMA co-occur with each other at all spatial scales.

LMA represents the tradeoff between fast growth and leaf construction costs (and longer leaf lifespans) (van Bodegom et al., 2014; Westoby et al., 2000, Wright et al., 2002). Across scales, LMA reflects different strategies for resource acquisition and maintenance between cooccurring species (Poorter et al., 2009), and within a species in different environments (Asner et al., 2011). As a consequence, variation in LMA is influenced by multiple factors. Field studies generally concur that inter-species differences are the single greatest source of variation in LMA. For example, in a study of hundreds of species across China, Yang et al. (2019) showed that climate explained only 15% of the variation in specific leaf area (SLA, the reciprocal of LMA), while taxonomic family explained 40%. As well, LMA varies significantly between fully sunlit and shaded leaves (Ellsworth & Reich, 1993; Keenan & Niinemets, 2016; Niinemets et al., 2015), which can be difficult to disentangle using remote sensing.

### 5.3 Implications and shortcomings

The unique location, complex terrain, and dynamic climates create diverse forests in the Western Ghats, but sparse field data poses great challenge in studying its plant functional traits and trait-environment relationships. Our study highlighted the capability of imaging spectroscopy in retrieving trait information in such regions. Trait maps from our study could fill some gaps in the field data and databases as noted by Jetz et al. (2016). As well, trait maps provide spatially continuous trait data for large areas and make it possible to conduct studies at different scales.

Our study revealed the preservation of trait space during the aggregation from plot level to regional level. It implies that the trade-offs among different plant functional traits can also be studied using trait maps with coarser resolutions, which underlies the huge potential of space-borne hyperspectral sensors in understanding ecosystem function.

For logistical reasons, we were unable to collect field samples concurrent with the AVIRIS flights. We took extra care to select plots without known disturbances after the image acquisition, but this mismatch still limited our study to traits that are relatively stable for a given phenological stage and possibly affected our trait model performances. Nevertheless, other studies in natural ecosystems with data collected across multiple years show relative stability in trait predictions across years, again provided that the seasonal timing and climatic conditions are comparable (Wang et al., 2020). We also should note that due to cloud cover our imagery was collected during the dry season, during which some deciduous species would have shed their leaves and thus be excluded from our analyses. We would expect greater trait variation and a potentially more diverse trait space for the Western Ghats during the wet season.

## Conclusion

Imaging spectroscopy enabled us to map seven foliar functional traits across six sites spanning the Western Ghats of India, which was not possible with existing in-situ trait data. Using the trait maps, we assessed trait-trait relationships as well as potential environmental drivers of trait distributions.

The traits exhibited similar distributions at each site. Distributions were generally unimodal, except for nitrogen concentration in Mudumalai where tea plantations contributed to a second peak around 30 mg g^−1^ compared to the median N of 20 mg g^−1^ elsewhere at Mudumalai and at other sites. Principal components analyses of trait estimates conducted at a range of data aggregation scales from native 4m to 1000m revealed a consistent pattern of trait space, with one axis representing the traditional leaf economic spectrum defined by nitrogen and LMA and another axis representing leaf structure and defense defined by fiber, lignin, and total phenolics. Multi-scale analysis of environmental drivers of traits showed that predictive models performed better at coarser resolutions for most traits, revealing strong broad-scale environmental controls on general trait patterns. The environmental variables did not capture the relevant local scale environmental drivers, or patterns may have been driven by other factors such as fine-scale soil variation, disturbance and land use history, competition, or ontogeny. Our multiscale analysis of trait variation from imaging spectroscopy demonstrates that there are broadscale drivers of trait variation that emerge at all scales. Further work is necessary to understand drivers of fine-scale variation, which may require environmental data that are not available for all sites or are not spatially explicit.

Total phenolics, fiber, and lignin exhibited strong environmental dependencies, while variation in calcium, sugars, and nitrogen was strongly controlled by site, indicating potential environmental variation not represented in our data (e.g., as with calcium and soils/geology). Among all the traits, LMA was poorly predicted by our models, which is unsurprising since species identity is the main controlling factor on variation in LMA (Poorter et al. 2009), and species with differing values of LMA fulfill different ecological roles (e.g., through niche partitioning) across spatial scales.

The trait data derived from hyperspectral imagery should provide additional ecological insights into ecosystem functioning in the Western Ghats. These data may provide suitable surrogates for diversity (i.e., using functional diversity, Asner et al., 2014; Schneider et al., 2017) or could provide essential information to parameterize ecosystem models (Butler et al., 2017; Van Bodegom et al., 2012), but in absence of spatially explicit species composition data, will not substitute for species based analyses derived from traditionally collected leaf-level trait data and databases.

Hyperspectral remote sensing on contemporaneous spaceborne platforms such as PRISMA (Italian Space Agency, ASI) EnMap (German Aerospace Centre, DLR) and HISUI (Japan Space Systems) and forthcoming missions like CHIME and SBG will expand our ability to retrieve plant functional traits globally and better understand drivers of trait variability and understand how traits may be changing as a consequence of climate change, especially in areas like the Western Ghats that are under-represented by in-situ sampling. The resulting data will enable comprehensively testing relationships across biomes and environmental gradients with much larger data sets or more spatial coverage than synthesis studies from databases.

## Acknowledgments

Funding for this research was provided by NASA grant 80NSSC17K0677 to PAT, AS and ARD. The authors gratefully acknowledge the assistance with fieldwork and logistics provided by Anish Sadanand, SynopticSense Inc., as well as the investment of the Indian Space Research Organisation (ISRO) and NASA in the collaborative data collection in India.

