## Supplemental Materials for "Variability in Forest Plant Traits along the Western Ghats of India and Their Environmental Drivers at Different Resolutions"

**S1 Filter plot spectra using PCA**

We applied the principal component analysis on the extracted spectra for each plot. The central point for the spectra from the same plot was defined as (PC1_mean_, PC2_mean_) where:

$${PC1}_{mean}=\frac{\sum_{i=1}^{n} {PC1}_{i}}{n}$$

$${PC2}_{mean}=\frac{\sum_{i=1}^{n} {PC2}_{i}}{n}$$

Where PC1_i_ (PC2_i_) is the first (second) component for the ith spectrum; n is the total number of spectra in the same plot. PC1_mean_ (PC2_mean_) is the mean of the first (second) component for all the spectra of the same plot.

The Euclidean distance of each spectrum to the central point was calculated following:

$$D = \sqrt{{({PC1}_{i}-{PC1}_{mean})}^{2}+{({PC2}_{i}-{PC2}_{mean})}^{2}}$$

We then calculated the mean and standard deviation of the distance. Spectra with D bigger than D_mean_+1.25D_sd_ or smaller than D_mean_ - 1.25D_sd_ were dropped.

**S2 Mask the shadowed pixels based on the brightness in near-infrared (NIR) region**

To mask the shadowed pixels on an image, we first calculated the norm of the NIR bands ([1017.55nm,1097.69nm]) for all the pixels. Secondly, we calculated the mean and the standard deviation of the NIR norm for the image. Pixels with NIR norm smaller than Norm_mean_ – 1.5Norm_sd_ were marked as shadows.

Table S1. Summary of PLSR models built using dry leaf spectra to estimate foliar traits. R^2^ and NRMSE are validation results based on withheld testing data.

| **Trait** | **Unit** | **Number of samples** | **Min** | **Max** | **R^2^** | **NRMSE** |
| --- | --- | --- | --- | --- | --- | --- |
| Calcium | mg/g | 541 | 0.5 | 64.1 | 0.76 | 10.04 |
| Fiber | mg/g | 758 | 86.2 | 731.2 | 0.78 | 10.83 |
| Lignin | mg/g | 768 | 5.8 | 438.2 | 0.84 | 10.49 |
| Nitrogen | mg/g | 1138 | 4.8 | 56.3 | 0.95 | 5.03 |
| Sugar | mg/g | 696 | 41.2 | 449.5 | 0.64 | 12.68 |
| Total Phenolics | mg/g | 635 | 12.6 | 316 | 0.77 | 9.00 |

Table S2. Variance inflation factor for the predictors at different resolutions. After removing elevation, the VIFs for all predictors are smaller than 10.

|  | **Elevation** | **MAT** | **MAP** | **Radiation** | **Seasonality** | **TWI** |
| --- | --- | --- | --- | --- | --- | --- |
| 30m | 34.75 | 37.70 | 7.38 | 8.41 | 5.20 | 1.06 |
| 100m | 34.12 | 36.53 | 7.18 | 8.04 | 4.83 | 1.12 |
| 1000m | 38.32 | 39.43 | 6.98 | 9.01 | 4.98 | 1.35 |
| 30m |  | 3.76 | 5.67 | 8.01 | 5.03 | 1.02 |
| 100m |  | 3.53 | 5.60 | 7.70 | 4.69 | 1.04 |
| 1000m |  | 4.18 | 5.41 | 8.72 | 4.87 | 1.18 |

| Trait | Trait Range | Model | | Calibration | | | Validation | | |
| --- | --- | --- | --- | --- | --- | --- | --- | --- | --- |
|  |  | Wavelengths | γ | R^2^ | RMSE | NRMSE | R^2^ | RMSE | NRMSE |
| Fiber  (mg/g) | 233.59-640.92 | 400-2400 | 5 | 0.5 | 73.9 | 18.72 | 0.43 | 84.6 | 21.46 |
|  |  | 1000-2400 | 5 | 0.66 | 55.8 | 14.65 | 0.43 | 84.5 | 21.44 |
|  |  | 1400-2400 | 3 | 0.31 | 78.3 | 21.56 | 0.30 | 87.4 | 25.16 |
| N  (mg/g) | 7.81-45.99 | 400-2400 | 4 | 0.49 | 6.0 | 14.36 | 0.39 | 6.9 | 21.74 |
|  |  | 1000-2400 | 4 | 0.54 | 4.7 | 13.72 | 0.50 | 5.6 | 21.02 |
|  |  | 1400-2400 | 3 | 0.31 | 5.9 | 16.76 | 0.30 | 6.2 | 23.98 |
| Lignin  (mg/g) | 74.73-428.43 | 400-2400 | 3 | 0.52 | 56.0 | 17.8 | 0.37 | 66.8 | 20.97 |
|  |  | 1000-2400 | 3 | 0.54 | 56.1 | 17.30 | 0.45 | 60.5 | 18.84 |
|  |  | 1400-2400 | 1 | 0.34 | 67.5 | 21.11 | 0.31 | 66.1 | 22.6 |

Table S3. PLSR model performances using different wavelength regions for nitrogen, fiber, and lignin

| **Trait** | **Resolution(m)** | **Radiation** | **MAP** | **Seasonality** | **MAT** | **TWI** | **Autocovariate** |
| --- | --- | --- | --- | --- | --- | --- | --- |
| Sugar | 30 | 7.16% | 14.10% | 37.52% | 29.18% | 1.85% | 10.18% |
| Sugar | 100 | 7.68% | 16.57% | 35.49% | 27.11% | 4.36% | 8.80% |
| Sugar | 1000 | 5.63% | 2.25% | 12.03% | 48.69% | 1.02% | 30.38% |
| ln(Phenolics) | 30 | 20.16% | 29.95% | 8.38% | 28.05% | 0.68% | 12.79% |
| ln(Phenolics) | 100 | 16.45% | 28.23% | 10.31% | 31.63% | 1.95% | 11.43% |
| ln(Phenolics) | 1000 | 8.29% | 16.14% | 25.46% | 44.74% | 1.77% | 3.61% |
| Ca | 30 | 12.53% | 22.01% | 28.55% | 9.55% | 0.98% | 26.38% |
| Ca | 100 | 11.23% | 21.34% | 30.60% | 9.39% | 2.93% | 24.51% |
| Ca | 1000 | 9.69% | 18.43% | 26.99% | 9.09% | 15.99% | 19.81% |
| N | 30 | 3.98% | 8.12% | 17.32% | 63.04% | 0.50% | 7.04% |
| N | 100 | 3.52% | 6.28% | 17.29% | 60.45% | 0.31% | 12.15% |
| N | 1000 | 7.16% | 20.82% | 10.42% | 42.93% | 8.87% | 9.79% |
| LMA | 30 | 8.04% | 7.69% | 11.55% | 37.59% | 0.12% | 35.00% |
| LMA | 100 | 11.62% | 9.85% | 10.34% | 38.34% | 0.32% | 29.52% |
| LMA | 1000 | 13.14% | 9.89% | 11.37% | 39.55% | 0.81% | 25.23% |
| Fiber | 30 | 35.93% | 21.61% | 16.25% | 17.00% | 0.13% | 9.08% |
| Fiber | 100 | 37.95% | 19.87% | 14.01% | 17.73% | 0.25% | 10.18% |
| Fiber | 1000 | 45.83% | 20.75% | 7.45% | 16.18% | 1.99% | 7.80% |
| Lignin | 30 | 39.59% | 34.22% | 9.79% | 9.04% | 0.15% | 7.20% |
| Lignin | 100 | 40.69% | 31.12% | 9.62% | 10.85% | 1.07% | 6.65% |
| Lignin | 1000 | 44.98% | 31.88% | 9.65% | 9.65% | 1.36% | 2.48% |

Table S4. Relative contribution to model R^2^ for each predictor at different scales. Numbers in red indicate the most important predictors. Contributions from all predictors add up to 100%.

| **Site** | **Sugar** | **Phenolics** | **Ca** | **N** | **LMA** | **Fiber** | **Lignin** | **resolution** |
| --- | --- | --- | --- | --- | --- | --- | --- | --- |
| Mudumalai | 33.09% | 27.26% | 18.89% | 71.93% | 39.13% | 43.30% | 73.75% | 30m |
|  | 24.70% | 20.31% | 12.86% | 63.00% | 27.96% | 32.13% | 63.10% | 100m |
|  | 13.34% | 11.69% | 8.66% | 54.77% | 16.81% | 16.38% | 53.08% | 1000m |
| Muddur | 28.30% | 13.38% | 34.64% | 63.75% | 37.90% | 43.27% | 43.01% | 30m |
|  | 24.05% | 10.26% | 28.33% | 53.05% | 29.63% | 35.64% | 33.85% | 100m |
|  | 17.62% | 5.60% | 16.88% | 44.40% | 23.12% | 23.75% | 25.47% | 1000m |
| Shimoga | 41.52% | 45.66% | 43.27% | 76.38% | 47.06% | 51.76% | 89.39% | 30m |
|  | 31.39% | 37.93% | 35.34% | 66.76% | 36.99% | 44.22% | 79.54% | 100m |
|  | 18.15% | 26.49% | 26.68% | 45.21% | 20.65% | 27.51% | 59.06% | 1000m |
| Sholayar | 32.26% | 41.65% | 19.97% | 54.41% | 44.22% | 41.86% | 57.57% | 30m |
|  | 23.73% | 35.23% | 12.72% | 42.22% | 33.05% | 30.80% | 47.94% | 100m |
|  | 13.80% | 27.28% | 6.71% | 22.78% | 20.73% | 16.97% | 36.44% | 1000m |
| Shoolpaneshwar | 27.68% | 24.42% | 28.68% | 46.11% | 38.13% | 32.65% | 54.61% | 30m |
|  | 16.78% | 14.08% | 19.02% | 29.71% | 28.20% | 19.59% | 37.42% | 100m |
|  | 7.29% | 0.16% | 8.91% | 14.77% | 18.10% | 6.90% | 14.47% | 1000m |
| Vansda | 35.34% | 23.74% | 27.40% | 51.39% | 36.13% | 33.59% | 60.42% | 30m |
|  | 25.97% | 16.00% | 21.80% | 33.77% | 26.52% | 21.69% | 50.84% | 100m |
|  | 8.83% | 6.64% | 12.81% | 16.43% | 13.52% | 7.87% | 24.51% | 1000m |

Table S5. Trait variance of each site at different resolutions. Shown as percentage of the variance at 4m.

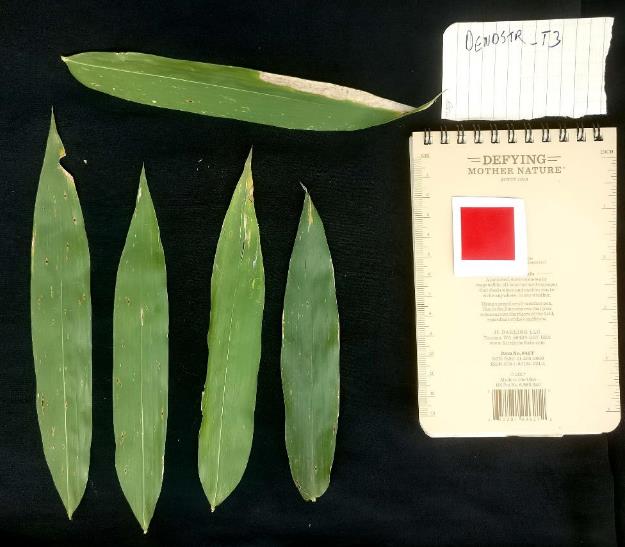
Fig. S1. Example photo of leaf area measurements. The red square is the reference object.

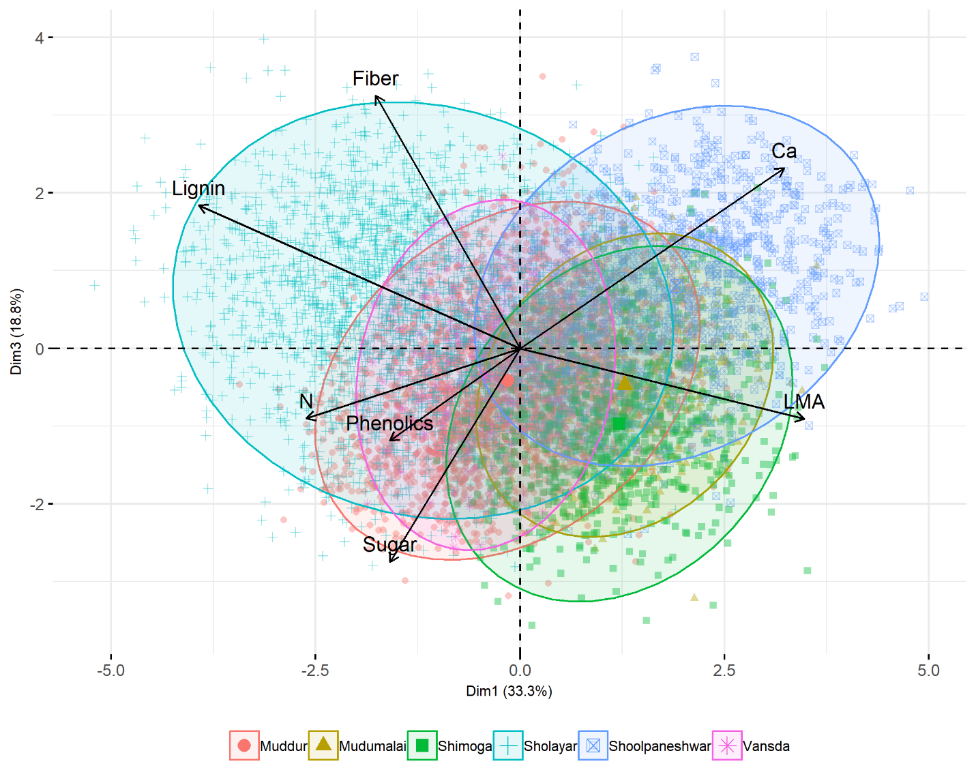

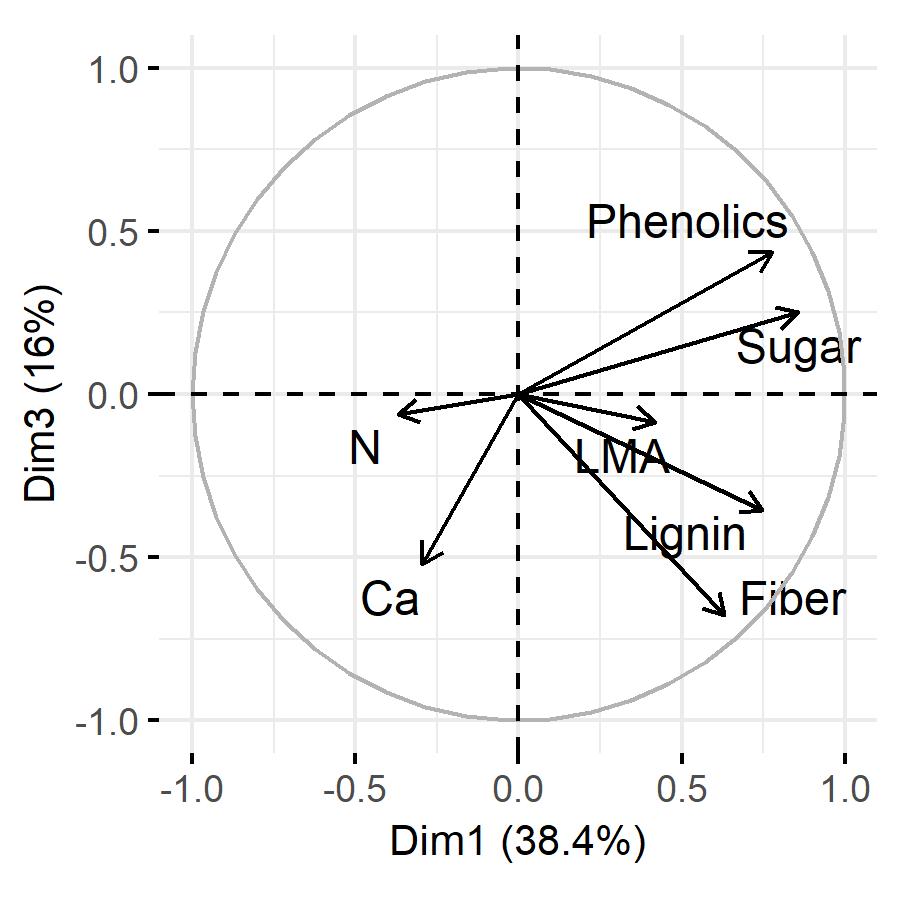

Fig. S2. Components 1 and 3 from PCA for image-level traits (~4m resolution) and the leaf-level traits (inset). Colored ellipses outline the trait space for different sites.

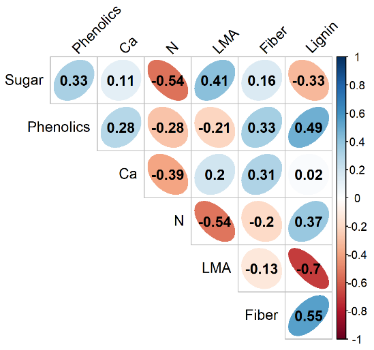

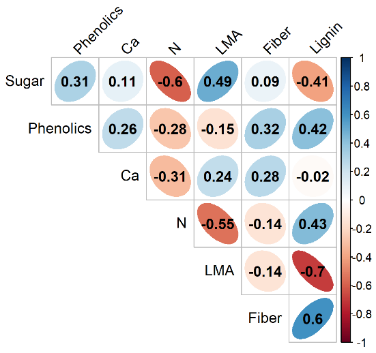

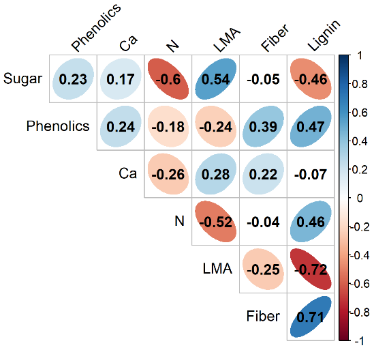

30m

100m

1000m

Fig. S3. Correlation between traits at three resolutions.

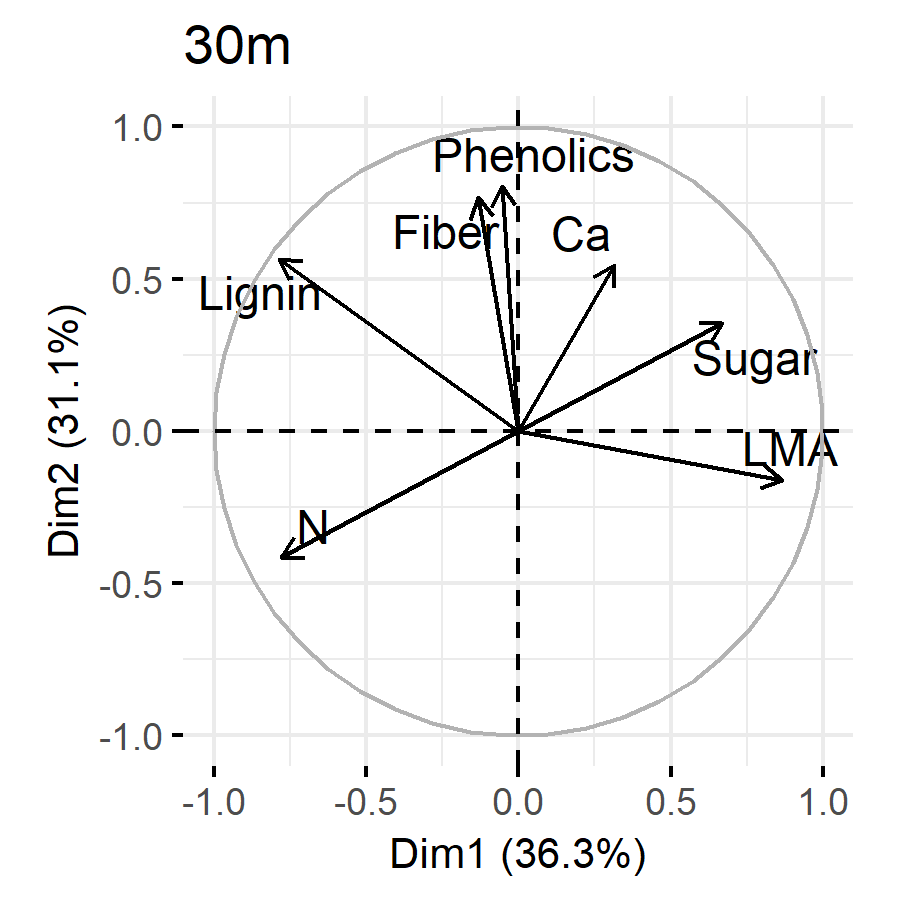

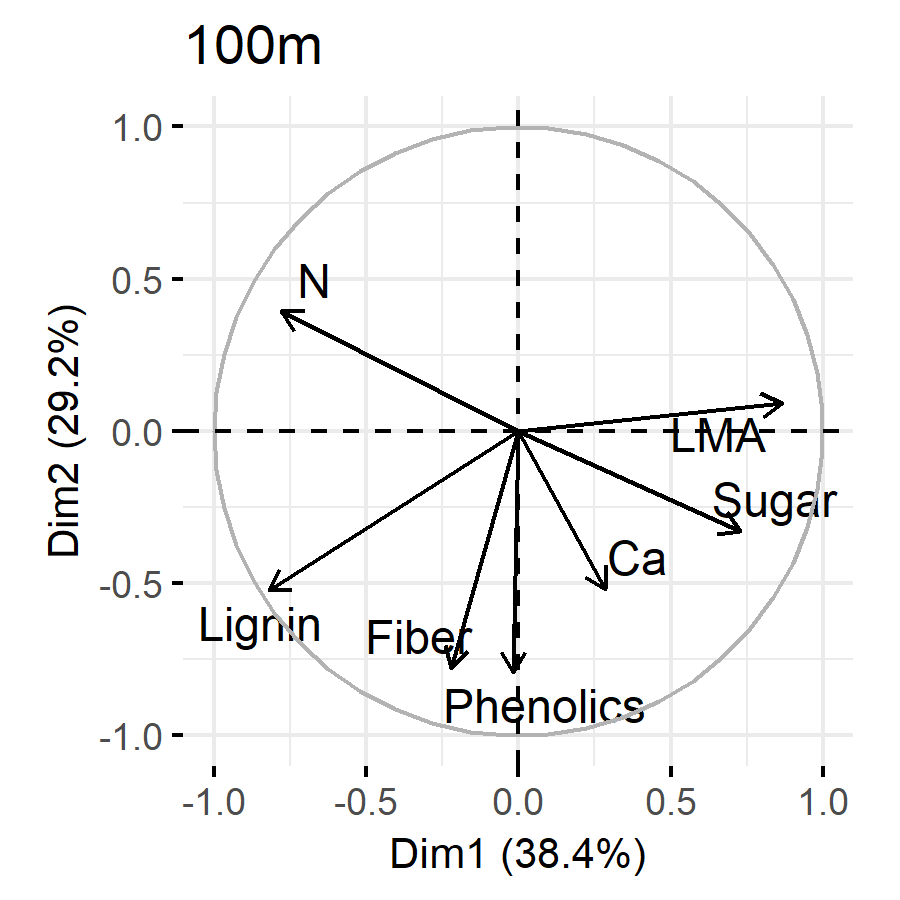

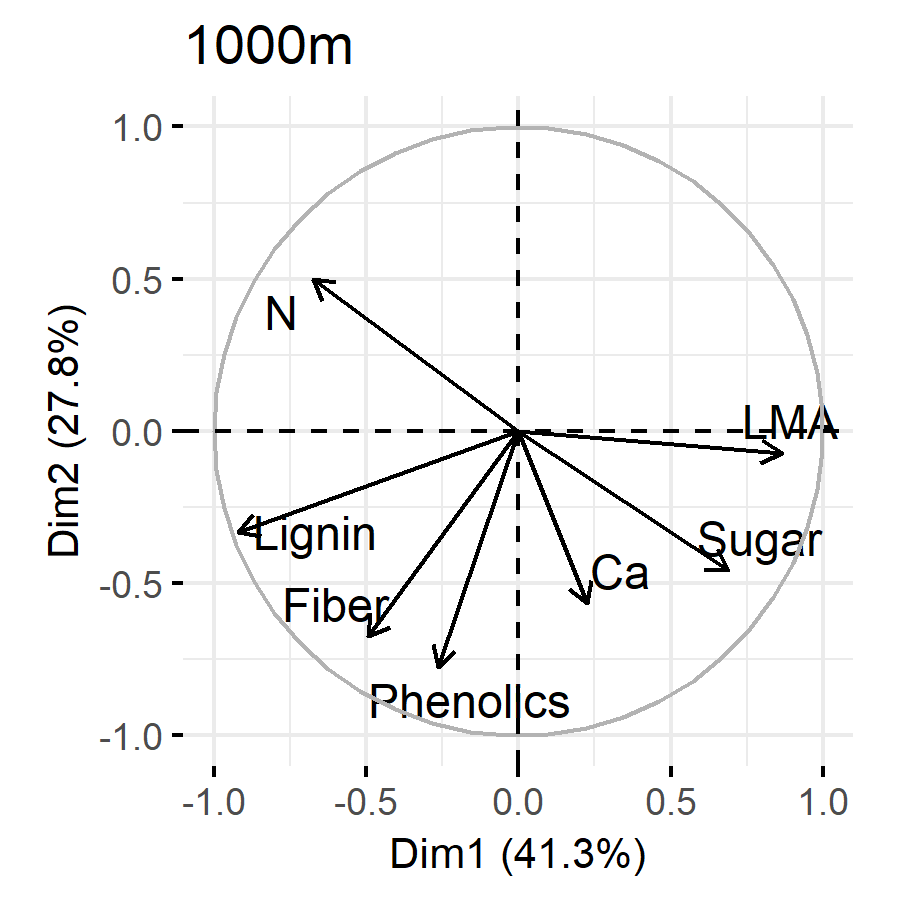

Fig. S4. PCA results for traits at three resolutions.

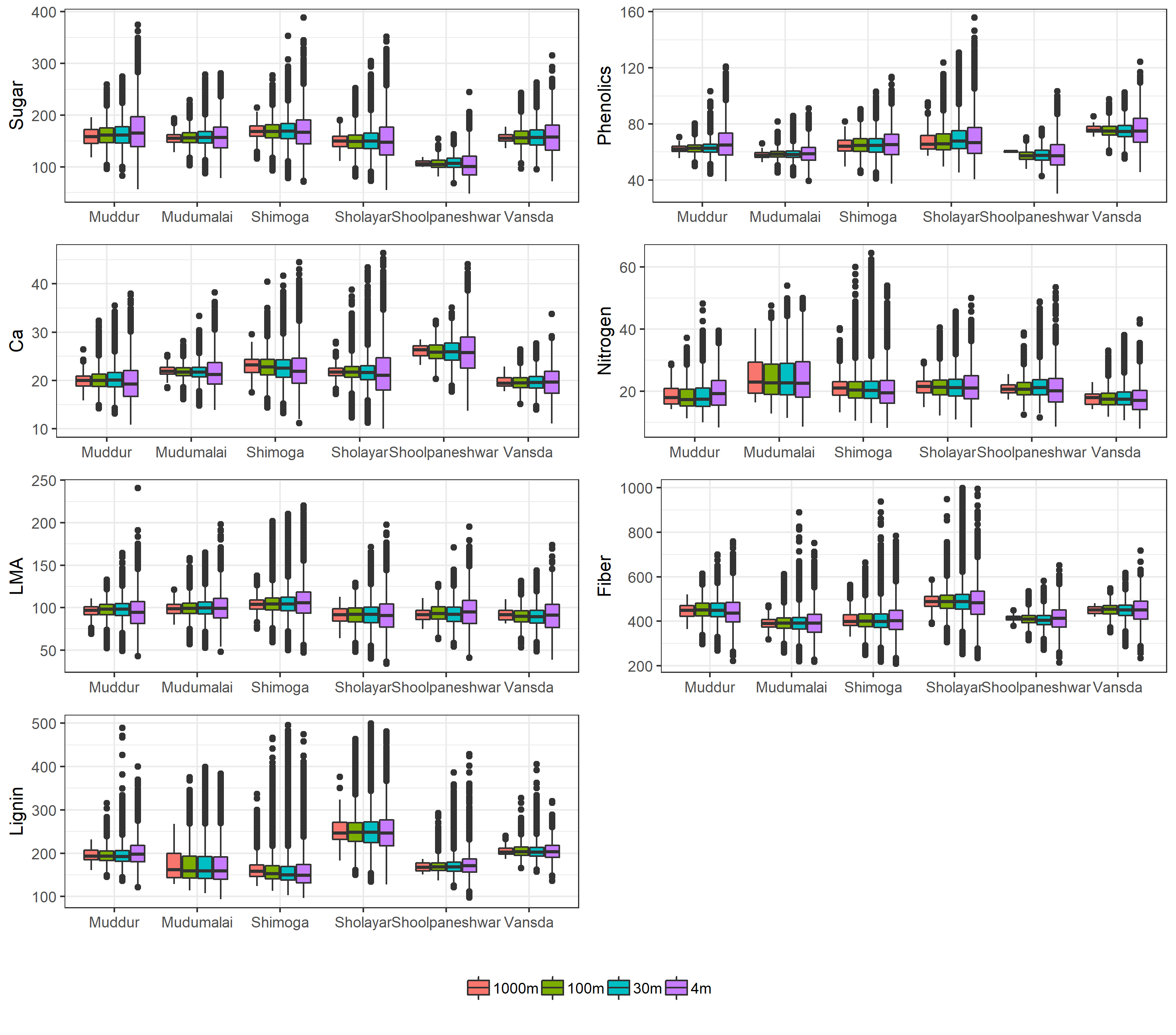

Fig. S5. Comparison of trait distribution across sites and resolutions.
